# TRPC6-Mediated Ca^2+^ Influx Activates MAPK and NFκB Signaling and Elicits Pro-Inflammatory and Catabolic Responses in Human Intervertebral Disc Cells

**DOI:** 10.64898/2026.02.07.704609

**Authors:** Janitri Venkatachala Babu, Varun Puvanesarajah, Addisu Mesfin, Jonathan P. Japa, Kevin Yoon, Mark Ehioghae, Michael G. Schrlau, Laura S. Stone, Wolfgang Hitzl, Karin Wuertz-Kozak

**Author notes:** Correspondence: Karin Wuertz-Kozak.

## Abstract

Intervertebral disc degeneration is characterized by inflammation, extracellular matrix breakdown, and neurovascular ingrowth, processes that contribute to discogenic, chronic back pain. The transient receptor potential canonical 6 (TRPC6) channel is a calcium-permeable ion channel implicated in inflammation and pain signaling in multiple tissues; however, its functional role in human disc cells remain unknown. Here, we investigated the expression, activation, and downstream consequences of TRPC6 activation using Hyp9, a pharmacological activator of TRPC6.

TRPC6 transcripts were consistently detected across all donors examined (n = 17). Functional TRPC6 activation induced a rapid, dose-dependent calcium (Ca^2+^) influx across 0.5–100 µM Hyp9. TRPC6 activation did not reduce metabolic activity or increase cytotoxicity at concentrations commonly used for in vitro TRPC6 activation. Mechanistically, TRPC6 activation induced mitogen-activated protein kinase (MAPK) and nuclear factor kappa B (NF-κB) pathways, as demonstrated by increased phosphorylation of p38 and extracellular signal-regulated kinase (ERK), degradation of the inhibitor of κB-alpha (IκB-α), and increased nuclear translocation of the NF-κB p65 subunit. Downstream of these early signaling events, TRPC6 activation elicited a robust inflammatory and catabolic response with upregulation of IL-6, IL-8, COX-2, MMP-1, MMP-3, NGF, and VEGF, with corresponding increases in protein secretion.

These findings identify TRPC6 as an important signaling node linking calcium influx to inflammatory, catabolic, and neuro- and angiogenesis-associated pathways in disc cells, highlighting TRPC6 as a potential therapeutic target in degenerative disc disease.

**Highlights:** *What are the main findings?:*

1. TRPC6 is endogenously expressed in human intervertebral disc cells, and its activation induces rapid calcium influx that initiates MAPK and NF-κB signaling pathways.
2. TRPC6 activation initiates a broad inflammatory and degenerative program, elevating the expression of IL-6, IL-8, COX-2, MMP-1, MMP-3, NGF, and VEGF.

*What are the implications of the main findings?:*

1. TRPC6 functions as a key upstream regulator linking calcium influx with inflammatory, matrix-degrading, and neuro-angiogenic processes central to disc degeneration and discogenic back pain.
2. Pharmacological targeting of TRPC6 may offer a novel therapeutic approach to suppress early inflammatory signaling, limit extracellular matrix breakdown, and reduce neurovascular ingrowth in degenerative disc disease.

## 1. Introduction

Low back pain is the leading cause of disability worldwide, with approximately 40% of cases attributed to discogenic etiology [1, 2]. Discogenic chronic back pain (DCBP), which arises from the degeneration of intervertebral discs (IVDs), is a multifactorial condition influenced by aging, genetic predisposition, mechanical overload, and environmental factors such as smoking and obesity [3-5]. The IVD is a fibrocartilaginous joint positioned between adjacent vertebral bodies composed of three primary components: the outer annulus fibrosus (AF), the central nucleus pulposus (NP), and the cartilaginous endplates that anchor the disc to the vertebrae [6]. Under healthy conditions, the disc is avascular, aneural, and largely immune-privileged, maintaining homeostasis through diffusion-dependent nutrient exchange and limited immune surveillance [7-9]. These unique characteristics help preserve tissue integrity but also limit the ability to address injury or inflammation once degeneration begins.

During degeneration, this finely tuned equilibrium of the disc is progressively disrupted [10]. Structural and biochemical deterioration manifests as a loss of proteoglycans, decreased hydration, breakdown of the extracellular matrix (ECM), and blurring of the AF-NP boundary. The resulting microenvironment is characterized by cellular stress, oxidative imbalance, and infiltration of immune and endothelial cells through breached endplates [11]. Aberrant vascular and neuronal ingrowth into the normally avascular and aneural tissue further exacerbates nociceptive sensitivity [12-14].

Together, these processes establish a chronic inflammatory-catabolic loop in which cytokines such as interleukin (IL)-6, IL-8, and tumor necrosis factor-α (TNF-α), as well as the prostaglandin-producing enzyme cyclooxygenase-2 (COX-2) and matrix-degrading enzymes including matrix metalloproteinases (MMPs) and a disintegrin and metalloproteinase with thrombospondin motifs (ADAMTS), accelerate ECM degradation and perpetuate pain [8]. Despite growing knowledge of these degenerative cascades, the upstream molecular sensors that convert mechanical and chemical stress into inflammatory signaling within the disc remain poorly characterized.

Among the putative molecular mediators of these pathological changes, the transient receptor potential canonical 6 (TRPC6) channel has emerged as a stress-responsive, calcium-permeable channel that integrates diverse physical and chemical stimuli [15]. TRPC6 is a diacylglycerol (DAG)-activated [16], non-selective cation channel that operates downstream of phospholipase C (PLC) and G-protein-coupled receptor (GPCR) signaling [17]. Its activity can also be modulated by factors such as hypo-osmotic conditions [18], membrane stretch [19], and the lipid mediator lysophosphatidylcholine (LysoPC) [20-22].

Under physiological conditions, TRPC6 contributes to the regulation of vascular tone [23], cytoskeletal remodeling [24, 25], and cellular proliferation [26]. In contrast, dysregulated TRPC6 activity has been associated with several pathological conditions, including renal fibrosis [27, 28], pulmonary hypertension and inflammation [29], cardiac hypertrophy [30], and neuroinflammation [31, 32]. Across these biological contexts, evidence supports the conclusion that TRPC6-mediated calcium influx activates various pathways, including NF-κB (nuclear factor kappa B) [33], MAPK (mitogen-activated protein kinase) [34], and RhoA (Ras homolog family member A) [35], driving the expression of pro-inflammatory, pro-fibrotic, and matrix-remodeling genes.

Beyond its established roles in inflammation and fibrosis in certain tissues, TRPC6 has also been implicated in pain signaling. It is expressed in peripheral sensory neurons, where its activation contributes to inflammatory hyperalgesia and neuropathic pain by modulating neuronal excitability and cytokine release [36-40]. Consequently, TRPC6 has gained increasing attention as a therapeutic target for inflammatory and pain-related disorders [41-43]. Despite this well-characterized involvement in various pathological conditions, the functional role of TRPC6 within the IVD, an immune-restricted, avascular tissue that is constantly exposed to high mechanical stress, remains poorly understood and particularly relevant to disc biology.

Previous studies, including our own [44-46], have demonstrated that TRPC6 is expressed in human IVD tissue, with higher levels detected in degenerated and painful discs. Within this context, TRPC6 may integrate polymodal cues, amplifying calcium-dependent activation of MMPs and cytokines that perpetuate degeneration. However, the mechanistic link between TRPC6 activation and inflammatory-catabolic signaling in disc cells remains undefined.

Given the emerging evidence linking TRPC6 activation to inflammatory and hyperalgesic responses across tissues, this study investigated whether pharmacological activation of TRPC6 induces inflammatory, catabolic, and neuro-angiogenic signaling programs in IVD cells. Using Hyp9, a TRPC6 pharmacological activator [47], we examined downstream responses of TRPC6 activation, including calcium influx, inflammatory and catabolic gene and protein expression, the regulation of neurotrophic and angiogenic mediators, and identification of involved pathways. These findings aim to establish TRPC6-associated calcium signaling as a potential early driver of key pathological cascades that underlie human disc degeneration and pain.

## 2. Materials and Methods

### 2.1 Human IVD Cell Isolation and Culture

IVD tissue was obtained from patients undergoing spinal surgery for disc herniation (DH) or degenerative disc disease (DDD) in accordance with institutional ethical guidelines (ethics approval number: URMC = STUDY00005200; Medstar = STUDY00008525). The mixed IVD tissue, comprising NP and AF regions, was excised intraoperatively, finely minced, and digested overnight at 37 °C and 5% CO_2_ in 1X Dulbecco’s Phosphate-Buffered Saline (DPBS; Cytiva, Marlborough, MA, USA; SH30028) containing 0.2% Collagenase NB4 (Nordmark Biochemicals, Uetersen, Germany; S1745401), 0.3% Dispase II (Roche, Mannheim, Germany; 04942078001), and 3% antibiotic-antimycotic (Cytiva, Marlborough, MA, USA; SV30079.01).

Due to the loss of distinct NP/AF boundaries in degenerated discs, isolating NP and AF cell types separately was not possible; therefore, these samples were categorized as a “mixed” IVD cell population representative of the overall disc microenvironment. Following enzymatic digestion, the cell suspension was filtered through a 70 µm cell strainer (EasyStrainer™, Greiner Bio-One; Kremsmünster, Austria; Cat. No. 542070), centrifuged at 300 × g for 10 minutes, resuspended in growth medium, and centrifuged again at 300 × g for 5 minutes prior to final resuspension. Cells were resuspended in growth medium consisting of HyClone DMEM/F12 1:1 medium with L-glutamine, HEPES (Cytiva, Marlborough, MA, USA; SH30023), supplemented with 10% fetal bovine serum (FBS; Cytiva, Marlborough, MA, USA; SH30396) and 1% antibiotic–antimycotic. Cells were maintained at 37 °C and 5% CO_2_ and passaged one to three times (P1-P3) before use in experiments.

Patient characteristics, including age, sex, disc level, and degeneration grade, are summarized in Supplementary Table 1.

### 2.2 TRPC6 Activation

To pharmacologically activate TRPC6, Hyp9 (MedChemExpress, Monmouth Junction, NJ, USA; HY-N10756), a TRPC6 pharmacological activator, was prepared as a 10 mM stock in dimethyl sulfoxide (DMSO; Sigma-Aldrich, St. Louis, MO, USA; D2650) and stored in aliquots at −80 °C. Stocks were diluted to 1 µM for treatment, maintaining a final DMSO concentration of ≤ 0.01% (v/v). A low, non-cytotoxic Hyp9 concentration was selected for downstream signaling and gene expression analyses to achieve controlled Ca^2+^ activation while minimizing non-specific effects.

IVD cells were seeded in different formats depending on the downstream assay. Cells were plated in 6-well plates (3 × 10^5^ cells/well) for gene expression, PicoGreen DNA quantification, ELISA, and Western blotting; in black-walled 96-well plates (0.1 × 10^5^ cells/well) for calcium flux and viability assays; and on 48-well plates (0.12 × 10^5^ cells/well) for immunofluorescence staining. Cells were cultured overnight in complete medium at 37 °C with 5% CO_2_, serum-starved for 2 h prior to treatment. Cells were subsequently treated with 1 µM Hyp9 for 18, 24, or 48 h under serum-free (0%) or serum-replete (10%) conditions, as specified for each downstream assay. Vehicle controls received equivalent DMSO volumes. All experiments were performed using at least three independent biological replicates.

### 2.3 Calcium Flux and Kinetic Analysis

Intracellular calcium influx following TRPC6 activation was assessed using the Fura-2 QBT™ Calcium Assay Kit (Molecular Devices, San Jose, CA, USA; R8197). IVD cells were seeded in black-walled, clear-bottom 96-well plates as described in Section 2.2. The following day, cells were loaded with Fura-2 QBT dye according to the manufacturer’s instructions and incubated for 1 h at 37 °C in phenol red-free, serum-free medium (Cytiva, Marlborough, MA, USA; SH30272). A feeder plate containing 10X concentrated activators or controls was prepared in parallel. Calcium flux was initiated by automated addition of activators (final 1X concentration) using a FlexStation 3 multi-mode microplate reader (Molecular Devices, San Jose, CA, USA). Fluorescence was recorded at 340 nm and 380 nm with an emission wavelength of 510 nm, and the 340/380 ratio was used to quantify real-time intracellular Ca^2+^ changes.

For kinetic analysis, recordings were acquired at 5-second intervals for 5 minutes. Baseline fluorescence (F_0_) was defined as the mean 340/380 ratio during the 30-second pre-stimulation period. The ΔF/F_0_ response was used to derive peak Ca^2+^ influx (maximum post-stimulation signal), the time-to-peak response, and the area under the curve (AUC), which reflects the total Ca^2+^ load. For concentration-response analysis, peak ΔF/F_0_ values were fitted using nonlinear regression in GraphPad Prism with a four-parameter logistic model in which the Top parameter (Emax) was constrained. Because responses did not reach a clear saturation plateau within the acquisition window, Emax was fixed to a value slightly above the maximal observed response (Top = 5; maximal observed ΔF/F_0_ ≈ 4) to stabilize curve fitting. Bottom and Hill slope parameters were estimated from the data, and half-maximal effective concentration (EC_50_) values are therefore reported as apparent estimates under the constrained Emax assumption.

### 2.4 Cell Viability and Cytotoxicity

Cell metabolic activity, an indicator of viability, was assessed with the PrestoBlue™ Cell Viability Reagent (Thermo Scientific, Waltham, MA, USA; P50201), and cytotoxicity was measured in parallel using the CyQUANT™ Lactate Dehydrogenase (LDH) Cytotoxicity Assay (Thermo Scientific, Waltham, MA, USA; C20301). After treatment with Hyp9 or vehicle controls (Section 2.2), cells were incubated with a 1:10 dilution of PrestoBlue in 1X DPBS for 4 h at 37 °C. Fluorescence was measured using a FlexStation 3 reader (Ex 560 nm/Em 590 nm). Conditioned media from cells treated for 18 h, 24 h, or 48 h were collected for LDH quantification according to the manufacturer’s instructions. Absorbance at 490 nm (LDH activity) and 680 nm (background) was recorded using the FlexStation 3. All assays were performed in technical duplicates across three independent biological replicates.

### 2.5 Gene Expression

Total RNA was extracted from IVD cells under serum-free conditions, as described in Section 2.2, for the experiments presented in the main figures, using TRIzol™ Reagent and Phasemaker™ Tubes Complete System (Thermo Scientific, Waltham, MA, USA; A33251) according to the manufacturer’s instructions. RNA quantity and purity were determined using a NanoPhotometer (Implen, Munich, Germany; N50). One microgram of RNA was reverse-transcribed using the High-Capacity cDNA Reverse Transcription Kit with RNase Inhibitor (Thermo Scientific, Waltham, MA, USA; 4374967). Quantitative PCR (qPCR) was performed using TaqMan™ Fast Advanced Master Mix (Thermo Scientific, Waltham, MA, USA; 4444963) and gene-specific TaqMan™ Expression Assays on a QuantStudio™ 3 Real-Time PCR System (Thermo Scientific, Waltham, MA, USA). Relative gene expression was calculated using the 2^−ΔΔCt^ method and normalized to the housekeeping gene YWHAZ and the respective vehicle control. YWHAZ was used as the reference gene based on its stable Ct values across donors, time points, and treatment conditions in human intervertebral disc cells; TBP and GAPDH were also evaluated but showed greater variability. Fold changes (2^−ΔΔCt^) were summarized as the arithmetic mean ± SEM and plotted on a logarithmic scale; individual donor values are shown. All reactions were performed in technical duplicates from independent biological replicates (n = 5-17 donors; exact n per experiment is reported in the corresponding figure legends). A complete list of TaqMan™ assays is provided in Supplementary Table 2. Gene expression analyses performed under serum-replete conditions are provided in the Supplementary Materials.

### 2.6 PicoGreen Assay

Cellular DNA content was quantified using the Quant-iT™ PicoGreen™ dsDNA Assay Kit (Thermo Scientific, Waltham, MA, USA; P11495) to normalize ELISA measurements. IVD cells were treated with 1 µM Hyp9 for 18, 24, or 48 h, under serum-replete conditions, as described in Section 2.2. At each time point, conditioned media from the same wells were collected for ELISA (Section 2.7). Cells were then washed once with 1× PBS and lysed in TE buffer (Invitrogen; Carlsbad, CA, USA; Cat. No. AM9849) supplemented with 0.1% Triton X-100 (Sigma-Aldrich; St. Louis, MO, USA; Cat. No. T8787). Lysates were processed for DNA quantification according to the manufacturer’s protocol. DNA concentrations were calculated from a standard curve and used to normalize secreted mediator levels. Corresponding DNA normalization and ELISA data obtained under serum-free conditions are provided in the Supplementary Materials.

### 2.7 Enzyme-Linked Immunosorbent Assay (ELISA)

Secreted inflammatory, catabolic, and neuro-angiogenic mediators were quantified from conditioned media collected after Hyp9 or vehicle treatment (Section 2.2), performed under serum-replete conditions for experiments presented in the main figures. Media were centrifuged to remove debris and analyzed using human DuoSet ELISA kits (R&D Systems, Minneapolis, MN, USA; DY008B) following the manufacturer’s instructions. The analytes included IL-6 (DY206), IL-8 (DY208), MMP-1 (DY901B), MMP-3 (DY513), NGF (DY256), and VEGF (DY293B). Absorbance was measured at 450 nm, with correction at 570 nm, using the FlexStation 3. Concentrations were calculated from standard curves, normalized to DNA content (section 2.6), and expressed as fold change relative to vehicle controls. All ELISA measurements were performed in technical duplicates using samples from five independent biological replicates.

### 2.8 Immunofluorescence

IVD cells were plated on 48 well plate and allowed to adhere overnight. The following day, cells were serum-starved and treated with Hyp9 or vehicle controls as described in Section 2.2, with staining performed at 15- or 30-minutes post-treatment. Cells were washed twice with cold 1X DPBS and fixed with 4% paraformaldehyde (Thermo Scientific, Waltham, MA, USA; J19943.K2) for 15 minutes at room temperature.

For NF-κB p65 staining, fixed cells were permeabilized with 0.1% Triton X-100 (Sigma-Aldrich, St. Louis, MO, USA; T8787), blocked with 5% donkey serum in 1X DPBS and incubated with primary NF-κB p65 (1:200, Cell Signaling Technology, Danvers, MA, USA; 8242) overnight at 4 °C. Cells were then incubated with Alexa Fluor® 488–conjugated donkey anti-rabbit secondary antibody (1:500; Abcam, Cambridge, UK; AB150073).

For TRPC6 staining, permeabilization was omitted. Cells were blocked as described above and incubated with primary TRPC6 antibody (1:50, Alomone Labs, Jerusalem, Israel; ACC-120) overnight at 4 °C, followed by Alexa Fluor® 488-conjugated donkey anti-rabbit secondary antibody (1:500; Abcam, AB150073). Primary and secondary antibodies were diluted in 1× DPBS containing 1% BSA and 0.1% Triton X-100, which may allow limited antibody access to membrane-associated intracellular compartments following fixation.

For both staining protocols, nuclei were counterstained with Hoechst 33342 (1 µg/mL, Thermo Scientific, 62249), and cells were maintained in 1× DPBS prior to imaging. Images were acquired using an Olympus IX-81 inverted microscope with CellSens v4.3 software (Olympus, Tokyo, Japan) and processed using FIJI/ImageJ (NIH, Bethesda, MD, USA).

### 2.9 Western Blot

IVD cells were treated with Hyp9 or vehicle controls as described in Section 2.2, and lysates were collected at 15 minutes and 30 minutes post-treatment. Cells were washed with cold 1X DPBS and lysed in RIPA buffer (Thermo Scientific, Waltham, MA, USA; 89901) containing 1X protease and phosphatase inhibitors (Thermo Scientific, Waltham, MA, USA; 78444). Protein concentrations were measured using the Pierce™ BCA Assay (Thermo Scientific, Waltham, MA, USA; 23225).

Equal amounts of protein (12–15 µg) were resolved by SDS–PAGE on 4–20% precast gels (Bio-Rad, 4561094) and transferred to nitrocellulose membranes (Bio-Rad, Hercules, CA, USA; 1704158).

Membranes were blocked with 5% BSA in TBS-T (0.1% Tween-20) for 1 h and incubated overnight at 4 °C with primary antibodies against phospho- and total p38, ERK, IκB-α and NF-κB IκB-α (Supplementary Table 3). After washing, membranes were incubated with fluorescent secondary antibodies (1:20,000; LI-COR Biosciences, Lincoln, NE, USA) for 1 h in the dark. Blots were scanned using visible and near-infrared (NIR) fluorescent detection in Odyssey XF imaging system (LI-COR Biosciences, Lincoln, NE, USA) and analyzed with the Empiria Studio Software (LI-COR Biosciences, Lincoln, NE, USA). Protein expression levels were normalized to housekeeping controls (GAPDH or α-tubulin) and reported as phospho/total ratios.

### 2.10 Statistical Analysis

All statistical analyses were performed using GraphPad Prism (version 10.6.1; GraphPad Software, San Diego, CA, USA). Data are presented as mean ± SEM, and statistical significance was set at p < 0.05. For gene expression and protein secretion (ELISA) data comparing two groups, differences were assessed using the nonparametric Mann-Whitney U test. Calcium influx dose-response data, consisting of repeated measures across Hyp9 concentrations within the same donor, were analyzed using the Friedman test followed by Dunn’s multiple comparisons test. Cell viability and cytotoxicity data across treatment conditions were analyzed using repeated-measures two-way ANOVA, with Hyp9 concentration and serum condition (0% vs. 10%) as factors, followed by Tukey’s post hoc test for multiple comparisons. Quantification of signaling pathway activation (e.g., Western blot densitometry and phospho-protein ratios) was analyzed using the Mann-Whitney U test. The number of biological replicates (independent human donors) and technical replicates is specified in the corresponding figure legends.

## 3. Results

### 3.1. Basal TRPC6 Expression In Human IVD Cells

TRPC6 is a diacylglycerol-activated, non-selective cation channel implicated in mechanotransduction, calcium signaling, and inflammatory responses in various tissues [48]. Although TRPC6 expression has been documented at the whole-disc level [44-46], its abundance in isolated IVD cells has not been systematically characterized. Because Hyp9-mediated TRPC6 activation forms the mechanistic foundation of this study, we first quantified endogenous TRPC6 expression in primary IVD cells.

TRPC6 transcripts were consistently detected across all donors examined (n=17; Pfirrmann degeneration grades III–V), with a mean ΔCt of 11.30 ± 0.47 (range: 7.68 – 15.31) relative to the reference gene YWHAZ (Figure 1A). These data indicate moderate but stable expression, reflecting the biological heterogeneity inherent to primary samples. Immunofluorescence confirmed TRPC6 protein localization, revealing punctate cytoplasmic staining with enrichment in perinuclear regions (Figure 1B), a pattern consistent with functional TRPC6 channels and intracellular trafficking described in other cell types [42].

**Figure 1.**
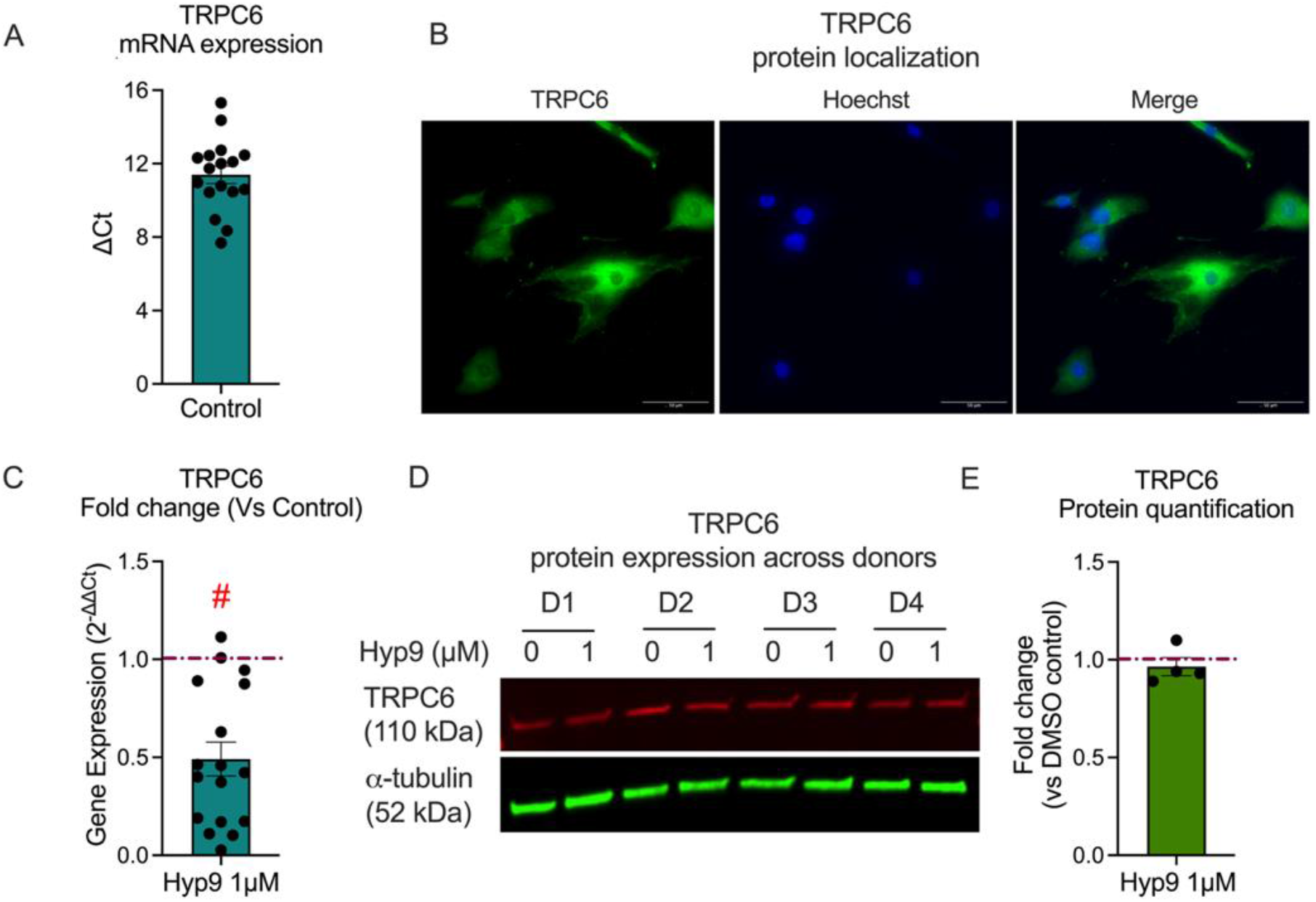
Endogenous expression of TRPC6 in human IVD cells. (A) Basal mRNA expression of TRPC6 in primary IVD cells (n=17), presented as ΔCt values normalized to YWHAZ. (B) Representative immunofluorescence staining of TRPC6 (green) in IVD cells, with punctate cytoplasmic staining reflecting functional channel pools and intracellular trafficking; nuclei counterstained with Hoechst 33342 (blue). Scale bar = 50 µm. (C) TRPC6 mRNA expression following 18 h treatment with Hyp9 (1 µM), expressed as fold change relative to vehicle control (red dashed line, n=17; Mann-Whitney test; #p < 0.0001). (D) Western blot analysis of TRPC6 (∼110 kDa) in cells treated with vehicle (0 µM) or Hyp9 (1 µM, 18 h); α-tubulin (52 kDa) serves as the loading control. (E) Densitometric quantification of TRPC6 protein levels across donors (n=4), presented as fold change relative to vehicle control (red dashed line, set to 1.0; Mann–Whitney test, p = 0.31). Data are presented as mean ± SEM; n denotes biological replicates.

To determine whether prolonged pharmacological activation alters TRPC6 transcript levels, primary IVD cells derived from degenerated discs were treated with Hyp9 (1 µM) for 18 h under serum-free conditions. TRPC6 mRNA expression was significantly reduced following Hyp9 treatment, with expression decreasing to 0.49 ± 0.07-fold relative to vehicle controls (Mann–Whitney test, p < 0.0001; Figure 1C), indicating a reduction in TRPC6 transcript levels following prolonged pharmacological activation under these experimental conditions.

To determine whether this transcriptional downregulation translated to altered protein abundance, TRPC6 protein levels were assessed following 18 h of Hyp9 treatment under serum-free conditions. Western blot analysis further validated TRPC6 expression, demonstrating a stable ∼110 kDa band corresponding to full-length TRPC6 in all donors tested (Figure 1D). Quantification showed that TRPC6 protein abundance remained unchanged following Hyp9 treatment, with a fold-change of 0.97 ± 0.05 (p = 0.31) relative to vehicle controls (Figure 1E). This indicates that short-term pharmacological activation does not acutely alter total TRPC6 protein levels, and the activator Hyp9 can be utilized in downstream assays without inducing compensatory changes in overall TRPC6 protein expression.

Collectively, these results demonstrate that IVD cells constitutively express TRPC6. Although sustained activation induces transcriptional downregulation of TRPC6 mRNA, TRPC6 protein stability is maintained during acute pharmacological activation, providing a foundation for subsequent Hyp9-mediated activation studies. Additional exploratory correlation analyses related to basal TRPC6 expression and donor-specific variables are provided in Supplementary Figure S1.

### 3.2. Hyp9-mediated TRPC6 Activation Induces Dose-dependent Calcium Influx and Altered Calcium Kinetics

Given that TRPC6 signaling relies on transmembrane cation influx, we next sought to confirm that the detected protein mediates functional calcium entry upon pharmacological activation. We assessed channel activity by monitoring real-time intracellular calcium kinetics ([Ca^2+^]_i_) using Fura-2-based ratiometric imaging.

Addition of Hyp9 elicited a rapid rise in intracellular Ca^2+^ in a concentration-dependent manner (Figure 2A). Peak ΔF/F_0_ values increased progressively from 1.12 ± 0.01 (0.5 µM) to 1.74 ± 0.15 (25 µM), 2.37 ± 0.21 (50 µM), and 3.26 ± 0.41 (100 µM). The kinetic profile exhibited an immediate rise in cytosolic calcium, followed by a sustained plateau, consistent with sustained channel opening that supports continuous cation entry rather than rapid desensitization.

**Figure 2.**
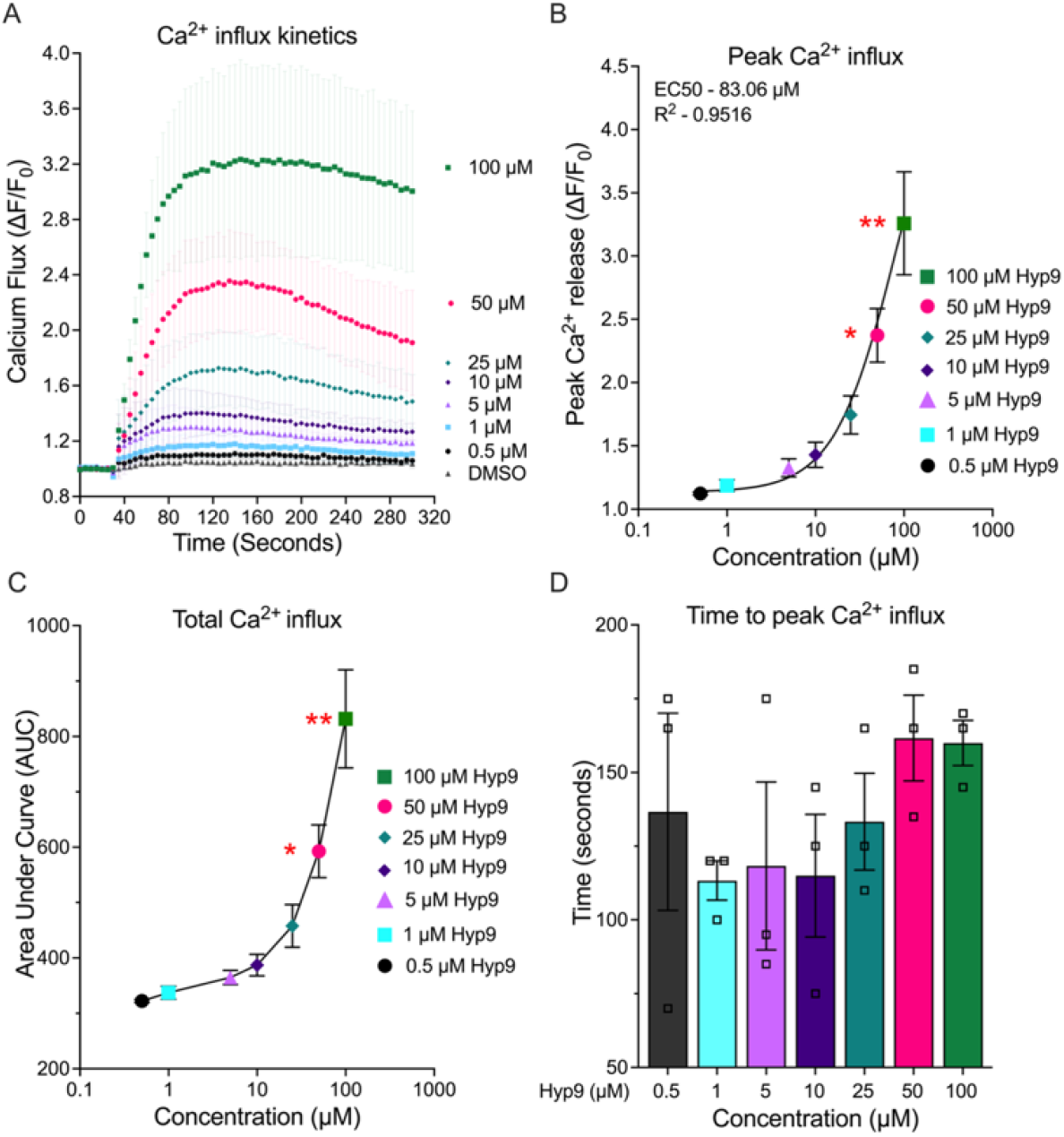
Hyp9-mediated TRPC6 activation induces dose-dependent Ca^2+^ influx in human IVD cells. (A) Real-time Ca^2+^ influx kinetics following acute Hyp9 stimulation (0.5–100 µM). (B) Peak Ca^2+^ influx (ΔF/F_0_) with nonlinear regression (EC_50_ = 83.06 µM; R^2^ = 0.95). (C) Total Ca^2+^ influx quantified as area under the curve (AUC). (D) Time to peak Ca^2+^ influx. All measurements were performed using the Fura-2 QBT assay on the FlexStation 3 platform, and values represent mean ± SEM (n = 3; biological replicates). Statistical significance (B-D) was assessed using Friedman test followed by Dunn’s multiple comparisons test, with comparisons made relative to DMSO control (*p < 0.05, **p < 0.01).

To define the potency and dynamic range of this response, nonlinear regression analysis was performed on the peak values. Concentration–response modeling yielded an EC_50_ of 83.06 µM (R^2^ = 0.95; Figure 2B). The relatively high EC_50_ observed for Hyp9 in primary IVD cells, compared with values reported in HEK293 systems overexpressing homomeric TRPC6, is consistent with the presence of native TRPC6-containing heterotetrameric channels (e.g., TRPC3/6/7 assemblies). Such heteromeric complexes would be expected to present fewer Hyp9 binding sites per channel than homotetrameric TRPC6, resulting in reduced apparent agonist potency in physiologic cells.

At lower concentrations (0.5–10 µM), Ca^2+^ increases were modest but reproducible, whereas concentrations ≥25 µM elicited steep, high-amplitude responses, consistent with the Hyp9-TRPC6 activation characteristics described in other systems [16]. Statistical analysis revealed that peak Ca^2+^ influx was significantly increased relative to DMSO controls with 50 µM (2.37 ± 0.21; p = 0.02) and 100 µM (3.26 ± 0.41; p = 0.003). Since peak values capture only a single time point, we also quantified the total calcium load using the Area Under the Curve (AUC). Total Ca^2+^ influx, quantified by AUC, followed a similar dose-responsive trajectory, rising from 322.40 ± 3.01 (0.5 µM) to 831.96 ± 88.73 (100 µM) (Figure 2C). Consistent with peak responses, AUC values were significantly elevated at 50 µM (p = 0.02) and 100 µM (p = 0.003) relative to DMSO. Interestingly, despite marked differences in amplitude, the time to peak Ca^2+^ influx remained largely consistent across doses (range: 113–162 seconds, Figure 2D). This temporal stability suggests that increasing Hyp9 concentrations does not significantly alter the kinetics of channel opening but rather modulates the magnitude of activation by recruiting a larger fraction of available channels. These results demonstrate that IVD cells possess physiologically active TRPC6 channels that respond to Hyp9 with predictable, dose-dependent kinetics, providing a functional basis for downstream signaling events.

### 3.3. Hyp9-mediated TRPC6 Activation Preserves Metabolic Activity and Cell Viability at low, non-cytotoxic Concentrations

Prolonged TRPC6 activation, accompanied by sustained higher intracellular calcium levels, can influence cellular stress responses and has been associated with altered cell viability, apoptosis, or cytotoxicity in various tissues, depending on cell type and stimulus intensity [49]. To ensure that downstream transcriptomic and signaling responses reflected TRPC6 activation rather than secondary cytotoxicity, we evaluated the effects of graded Hyp9 exposure on metabolic activity and membrane integrity using PrestoBlue and LDH assays under both serum-replete (10%) and serum-free (0%) conditions at 18, 24, and 48 h (Figure 3).

**Figure 3.**
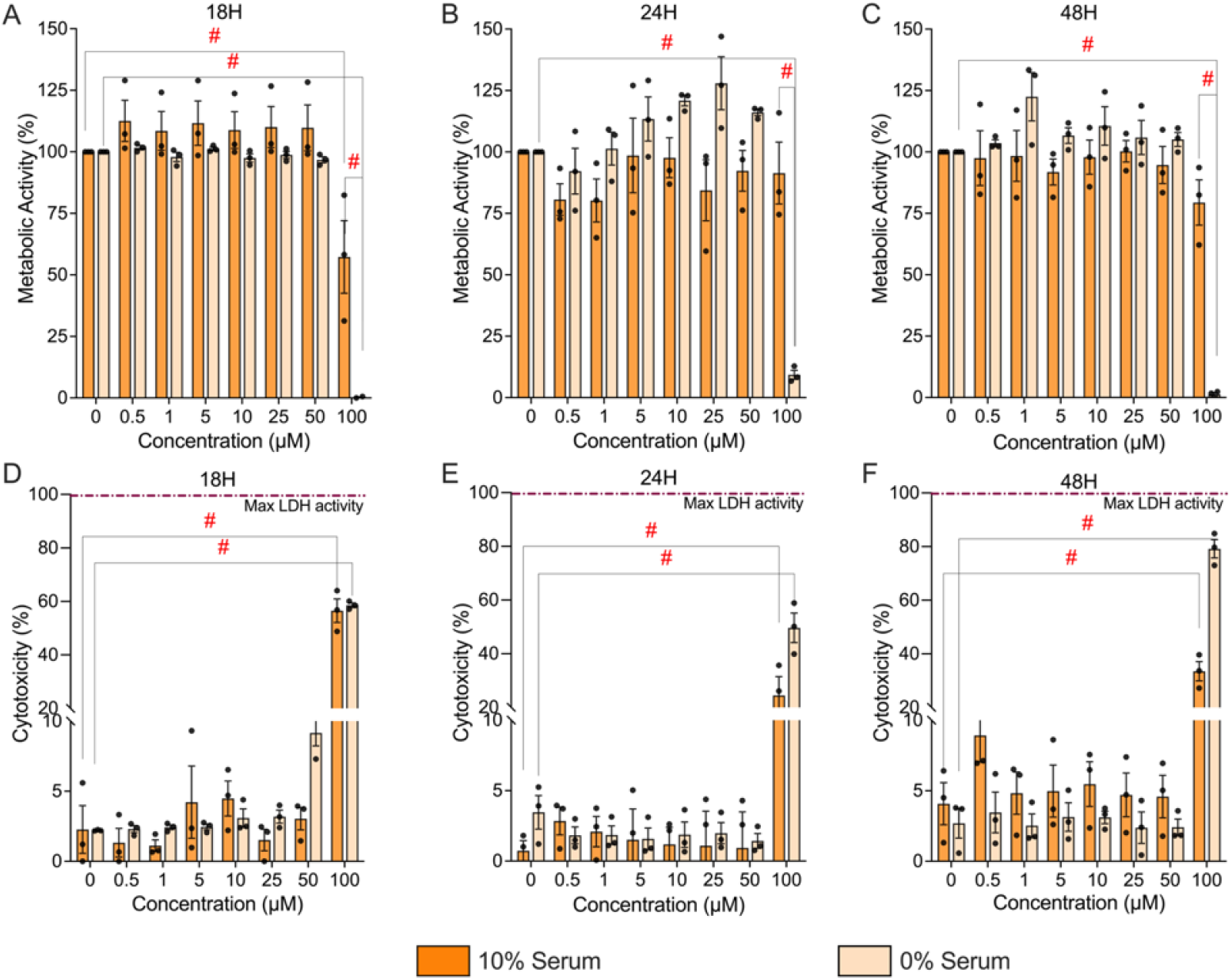
Concentration- and time-dependent effects of Hyp9 on metabolic activity and cytotoxicity in human IVD cells. (A-C) Metabolic activity assessed via the Presto Blue assay in IVD cells treated with Hyp9 (0–100 µM) for 18 h (A), 24 h (B), and 48 h (C). Data are stratified by culture conditions: 10% serum (orange) and 0% serum (beige). (D-F) Cytotoxicity quantified by LDH release after 18 h (D), 24 h (E), and 48 h (F) of exposure. The dashed magenta line indicates 100% maximal LDH activity (positive control). Data are presented as mean ± SEM; n=3, biological replicates. Statistical significance was assessed using two-way repeated-measures ANOVA (concentration × serum) followed by Tukey’s multiple comparisons test. (#p < 0.0001).

Across 10% serum conditions, metabolic activity remained stable up to 50 µM Hyp9 over 18–48 h (e.g., 50 µM at 18 h: 109.74 ± 9.30%, 24 h: 92.31 ± 8.30%, 48 h: 94.65 ± 7.55; Figure 3A–C). Under serum-free conditions, cell metabolism was similarly preserved at up to 50 µM Hyp9, with modest increases at 24 h (127.93 ± 10.78% for 25 µM) likely reflecting mild metabolic stimulation or donor variability. In contrast, the highest dose (100 µM) led to profound metabolic suppression in serum-free media (18 h: 0.19 ± 0.19%; p < 0.0001, 24 h: 9.31 ± 1.82%; p < 0.0001 48 h: 1.35 ± 0.75%; p < 0.0001) and partial reduction in 10% serum at early (18 h: 57.30 ± 14.74%; p < 0.0001) but not late timepoints..

To distinguish between metabolic suppression and actual cell death, we quantified cytotoxicity via LDH leakage. Consistent with the metabolic data, LDH release assays corroborated these findings (Figure 3D–F). Across 10% serum conditions, LDH levels remained low up to 50 µM Hyp9 for all time points (18 h: 3.05 ± 0.80%, 24 h: 0.94 ± 2.55%, 48 h: 4.59 ± 1.51%), indicating preserved membrane integrity even with prolonged exposure (Figure 3D–F). In contrast, 100 µM Hyp9 produced a pronounced cytotoxic response in 10% serum, increasing LDH release to 56.53 ± 4.39% (p < 0.0001) at 18 h and 33.51 ± 3.59% (p < 0.0001) at 48 h, with intermediate cytotoxicity at 24 h (24.47 ± 6.98%; p < 0.0001). Notably, serum-free conditions magnified the cytotoxic effect of Hyp9, consistent with the heightened sensitivity of IVD cells to metabolic stress in the absence of trophic support. At 50 µM Hyp9, LDH release was modest at 18 h (9.16 ± 0.93%) but decreased to near-baseline values at 24–48 h (1.44 ± 0.52%, 2.42 ± 0.58%), suggesting transient early stress without sustained membrane damage. However, 100 µM Hyp9 caused severe cytotoxicity in 0% serum: LDH release reached 58.57 ± 0.86% (p < 0.0001) at 18 h and escalated to 79.19 ± 3.42% (p < 0.0001) at 48 h, consistent with the profound metabolic collapse observed in the Presto Blue assays.

Taken together, these data demonstrate that IVD cells tolerate low, non-cytotoxic concentrations of Hyp9 (1 µM used in downstream experiments) and maintain metabolic and membrane integrity across extended exposure periods. In contrast, exposure to the highest concentration tested (100 µM) resulted in pronounced cytotoxicity, validating that the inflammatory, catabolic, and neurogenic responses observed in subsequent assays reflect TRPC6 activation rather than loss of cell viability.

### 3.4. Hyp9-mediated TRPC6 Activation Triggers a Robust Pro-inflammatory Response in Human IVD Cells

Dysregulated calcium signaling downstream of TRPC6 activation has been closely linked to inflammatory responses in multiple tissues [41], and pro-inflammatory mediators such as IL-6, IL-8, and COX-2 are strongly implicated in IVD degeneration and discogenic pain [50-52]. To determine whether TRPC6 activation contributes to a similar inflammatory program in IVD cells, we analyzed the transcriptional and translational expression of key inflammatory markers following Hyp9 stimulation.

Under serum-free conditions (0% serum), 18 h Hyp9 treatment (1 µM) resulted in a marked upregulation of all three inflammatory genes compared with vehicle (Figure 4A).

**Figure 4.**
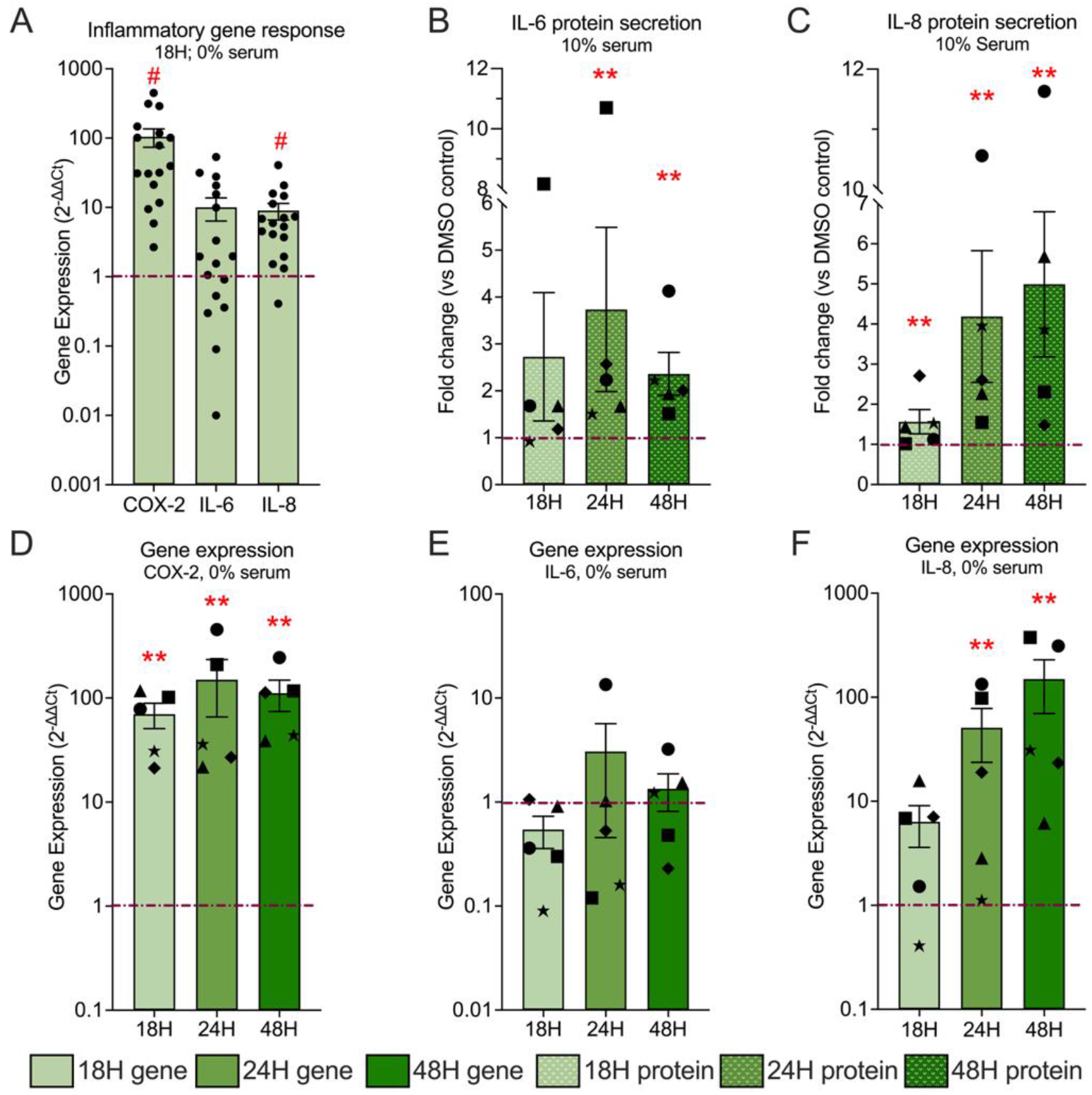
TRPC6 activation induces and sustains inflammatory gene expression and protein secretion in human IVD cells. (A) Transcriptional profiling of inflammatory markers after 18 h of Hyp9 (1 µM) treatment. COX-2, IL-6, and IL-8 mRNA levels are presented as fold change (2^−ΔΔCt^) relative to vehicle control (dashed line, n= 17). (B-C) Time-dependent secretion of IL-6 (B) and IL-8 (C) protein into the culture media, quantified by ELISA and normalized to total DNA content presented as fold change relative to vehicle control (dashed line, n=5). (D-F) Longitudinal gene expression analysis of COX-2 (D), IL-6 (E), and IL-8 (F) over 18, 24, and 48 h, demonstrating sustained transcriptional activation (n=5, symbols matched with protein secretion). Data are presented as mean ± SEM, with individual donor points; n denotes biological replicates. Statistical significance (**p<0.01, #p<0.0001 vs. vehicle control) was determined by the Mann-Whitney test compared to vehicle control.

IL-8 and COX-2 transcripts were significantly upregulated, with IL-8 increasing 9.01 ± 2.40-fold (p < 0.0001) and COX-2 exhibiting the most pronounced response, increasing 104.63 ± 31.08-fold (p < 0.0001; 2^−ΔΔCt^, n = 17 donors). IL-6 expression showed a trend toward induction (10.06 ± 3.68-fold; p = 0.12), reflecting substantial inter-donor variability. This indicates that while cytokine responses display biological heterogeneity, COX-2 upregulation is a highly conserved and uniform response to TRPC6 activation in IVD cells.

For a molecular mechanism to drive chronic disease, the response must be sustained rather than transient. To this end, we quantified COX-2, IL-6, and IL-8 expression under serum-free conditions across 18, 24, and 48 h in a smaller sample (Figure 4D–F, n=5). COX-2 (Figure 4D) displayed persistent, high-magnitude induction throughout all time points. Expression increased from 69.85 ± 18.99-fold (p = 0.008) at 18 h to 149.90 ± 84.0-fold (p = 0.008) at 24 h and remained strongly elevated at 111.2 ± 37.11-fold (p = 0.008) at 48 h. Importantly, sustained COX-2 upregulation was observed across the donor cohort, validating it as a robust and persistent downstream effector of TRPC6 signaling. IL-6 (Figure 4E) exhibited pronounced inter-donor variability across all time points.

Although mean expression values differed at 18, 24, and 48 h, none of these changes reached statistical significance (18 h: 0.54 ± 0.18-fold, p = 0.12; 24 h: 3.06 ± 2.60-fold, p = 0.68; 48 h: 1.34 ± 0.52-fold, p = 0.68), limiting the ability to draw definitive conclusions regarding temporal regulation of IL-6 transcription. IL-8 (Figure 4F) showed the greatest temporal dependence. Expression was already elevated at 18 h, albeit not significantly (6.33 ± 2.73-fold; p = 0.12), increased markedly at 24 h (50.87 ± 27.24-fold; p = 0.008), and remained high at 48 h (149.40 ± 79.85-fold; p = 0.008). Notably, IL-8 expression remained elevated above baseline in all donors at all time points, indicating consistent induction that was independent of quantitative variability.

To confirm that this transcriptional upregulation translated into a functional secretory phenotype, we quantified cytokine accumulation in the conditioned media with ELISA (Figure 4B-C). IL-6 and IL-8 release was measured under 10% serum conditions across 18, 24, and 48 h. IL-6 secretion increased to 2.72 ± 1.36-fold (p = 0.12) at 18 h and 3.73 ± 1.75-fold (p = 0.008) at 24 h before stabilizing at 2.36 ± 0.45-fold (p = 0.008) by 48 h. IL-8 secretion displayed a more gradual, sustained increase over time, rising from 1.56 ± 0.30-fold (18 h; p = 0.008) to 4.18 ± 1.6-fold (24 h; p = 0.008) and 4.99 ± 1.80-fold (48 h; p = 0.008). Supplementary analyses of inflammatory gene expression under serum-replete (10%) and cytokine secretion under serum-free (0%) conditions (Supplementary Figure S2) further confirm that the inflammatory phenotype is preserved across experimental contexts.

Taken together, the sustained transcriptional kinetics (Figure 4D-F) combined with progressively accumulating cytokine secretion (Figure 4B-C) demonstrate that TRPC6 activation is sufficient to elicit a robust, multi-phase pro-inflammatory program in IVD cells dominated by strong and persistent COX-2 induction and consistently elevated cytokine responses.

### 3.5. Hyp9-mediated TRPC6 Activation Selectively Regulates Collagenolytic and Aggrecan-Remodeling Pathways in Human IVD Cells

Loss of ECM integrity is a hallmark of disc degeneration, and MMPs are key mediators of proteoglycan and collagen breakdown in IVD tissue. They are strongly associated with progressive degeneration and pain [53, 54]. To determine whether TRPC6 activation contributes to catabolic ECM remodeling in IVD cells, we assessed the transcriptional regulation and protein secretion of key matrix-degrading enzymes following pharmacological activation with Hyp9 (1 µM).

Under serum-free conditions, acute Hyp9 treatment (18h) elicited a selective collagenolytic transcriptional response (Figure 5A). Among the MMPs examined, MMP-1 (collagenase-1) exhibited the strongest induction (20.03 ± 4.88-fold vs. vehicle; p < 0.0001), identifying it as a prominent early TRPC6-responsive gene. MMP-3 (stromelysin-1) showed a more modest but significant increase (2.33 ± 0.48-fold; p = 0.0003). In contrast, MMP-2 (gelatinase A) was significantly suppressed following Hyp9 treatment (0.81 ± 0.16-fold; p < 0.0001), while MMP-13 (collagenase-3) did not display a consistent transcriptional response (1.18 ± 0.18-fold; p = 0.36). These findings indicate that TRPC6 activation does not uniformly induce all collagenolytic enzymes but instead selectively recruits specific MMP family members.

**Figure 5.**
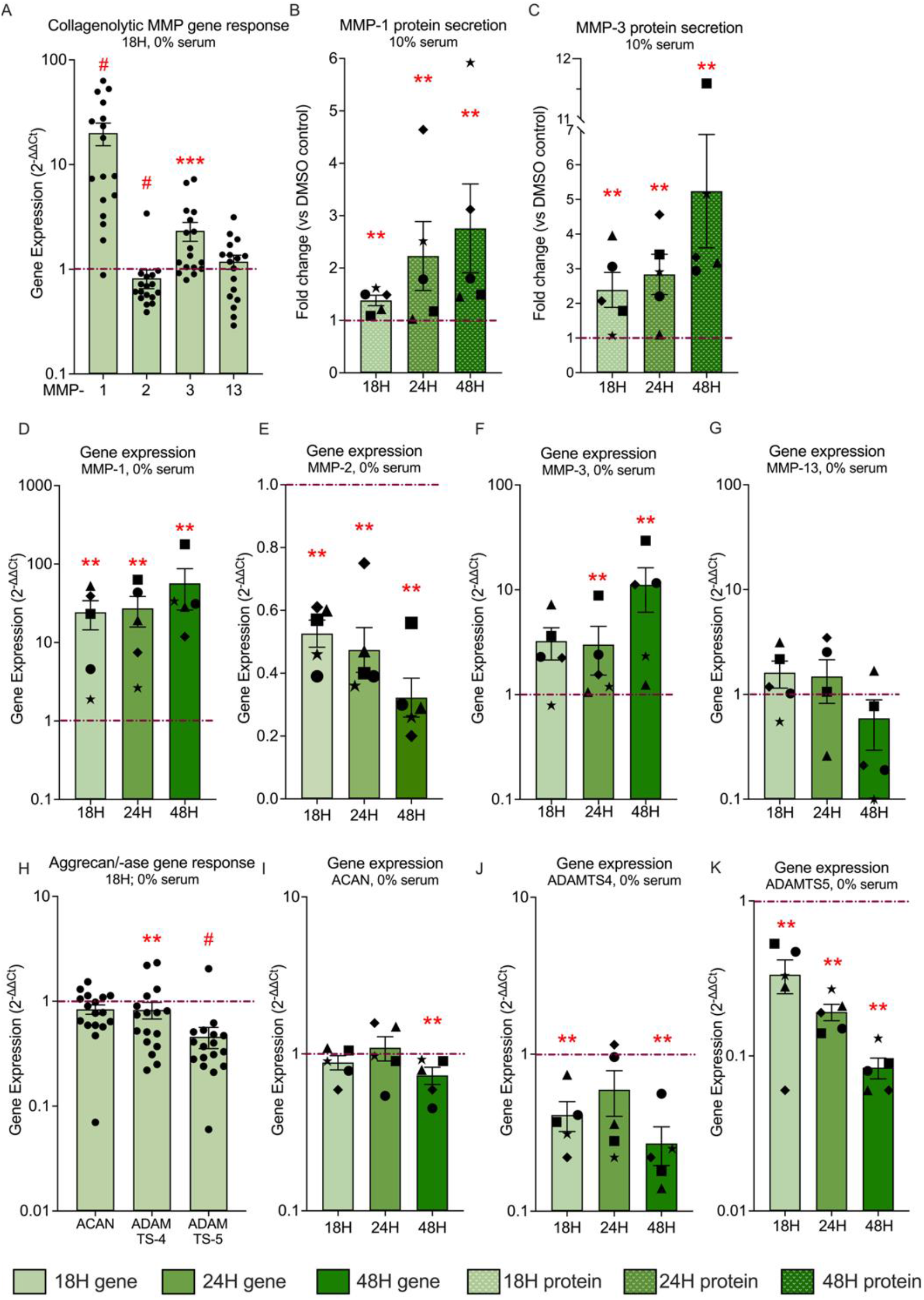
TRPC6 activation elicits selective collagenolytic and aggrecan-associated transcriptional remodeling in human IVD cells. (A) Transcriptional profiling of Collagenolytic MMP markers after 18 h of Hyp9 (1 µM) treatment. MMP-1, MMP-2, MMP-3, and MMP-13 mRNA levels are presented as fold change (2^−ΔΔCt^) relative to vehicle control (dashed line, n= 17). (B-C) Time-dependent secretion of MMP-1 (B) and MMP-3 (C) protein into the culture media, quantified by ELISA and normalized to total DNA content, presented as fold change relative to vehicle control (dashed line, n=5). (D-F) Longitudinal gene expression analysis of MMP-1 (D), MMP-2 (E) MMP-3 (F), and MMP-13 (G) over 18, 24, and 48 h, demonstrating sustained transcriptional activation. (Symbols represent matched biological donors across time points; n = 5). (H) Aggrecan (ACAN) and aggrecanase gene expression following 18 h of Hyp9 treatment under serum-free conditions, including ACAN, ADAMTS4, and ADAMTS5, presented as fold change relative to vehicle control (n = 17). (I–K) Temporal transcriptional regulation of ACAN (I), ADAMTS4 (J), and ADAMTS5 (K) under serum-free conditions following Hyp9 treatment at 18, 24, and 48 h (n = 5). Data are presented as mean ± SEM with individual donor points; n denotes biological replicates. Statistical significance (**p<0.01, ***p<0.001, #p<0.0001 vs. vehicle control) was determined by the Mann-Whitney test compared to vehicle control.

Time-course analysis under serum-free conditions further revealed distinct kinetic profiles among individual MMPs (Figure 5D–G). MMP-1 expression increased progressively over time, rising from 24.32 ± 9.80-fold at 18 h (p = 0.008) to 27.20 ± 11.43-fold at 24 h (p = 0.008) and reaching 56.72 ± 30.70-fold at 48 h (p = 0.008). This sustained and increasingly amplified response identifies MMP-1 as the primary catabolic target of TRPC6. MMP-3 exhibited a delayed induction pattern, with modest expression at early time points (3.24 ± 1.09-fold at 18 h; p = 0.12) followed by significant upregulation at 24 h (3.00 ± 1.50-fold; p = 0.008) and a pronounced increase at 48 h (11.15 ± 5.05-fold; p = 0.008). This is consistent with a delayed but robust induction pattern, suggesting a secondary wave of catabolic activation. In contrast, MMP-2 expression remained consistently and significantly suppressed across all time points, reaching 0.32 ± 0.08-fold by 48 h (p = 0.008), while MMP-13 expression showed inconsistent and non-significant regulation across donors at all time points, with mean expression values of 1.61 ± 0.46-fold at 18 h (p = 0.12), 1.48 ± 0.65-fold at 24 h (p = 0.68), and 0.60 ± 0.30-fold at 48 h (p = 0.12).

To determine whether transcriptional regulation translated into increased enzyme release, MMP protein secretion was quantified under serum-replete (10%) conditions to enable robust accumulation and detection of secreted proteins (Figure 5B–C). Hyp9 increased MMP-1 secretion from 1.38 ± 0.09-fold (18 h; p = 0.008) to 2.23 ± 0.65-fold (24 h; p = 0.008) and 2.76 ± 0.84-fold (48 h; p = 0.008), indicating progressive accumulation of MMP-1 protein over time. MMP-3 secretion followed a similar trajectory, rising from 2.38 ± 0.50-fold (18 h; p = 0.008) to 2.83 ± 0.57-fold (24 h; p = 0.008) and increasing further to 5.24 ± 1.63-fold (p = 0.008) by 48 h.

In parallel, analysis of aggrecan and aggrecanase/ADAMTS gene expression under serum-free conditions demonstrated pathway-specific and temporally distinct regulation (Figure 5H–K). At 18 h, ACAN expression was largely maintained (0.84 ± 0.08-fold; p = 0.12; n = 17), whereas ADAMTS4 (0.83 ± 0.15-fold; p = 0.008) and ADAMTS5 (0.45 ± 0.10-fold; p < 0.0001) were significantly suppressed relative to vehicle controls.

Extended temporal analysis demonstrated differential kinetics among aggrecan-associated genes. ACAN expression remained unchanged at 18 h (0.88 ± 0.09-fold; p = 0.68) and 24 h (1.09 ± 0.19-fold; p = 0.68), but showed a statistically significant, albeit minor, reduction at 48 h (0.73 ± 0.09-fold; p = 0.008). In contrast, ADAMTS4 was significantly suppressed at 18 h (0.41 ± 0.09-fold; p = 0.008), showed a similar but more variable reduction at 24 h that did not reach statistical significance (0.59 ± 0.19-fold; p = 0.13), and was again significantly reduced at 48 h (0.27 ± 0.07-fold; p = 0.008). ADAMTS5 expression was consistently and progressively suppressed across all time points, including 18 h (0.33 ± 0.08-fold; p = 0.008), 24 h (0.19 ± 0.02-fold; p = 0.008), and 48 h (0.08 ± 0.01-fold; p = 0.008). Supplementary analyses across varying serum conditions (Supplementary Figure S3) further supported these findings, including the observation that MMP-1 secretion remained robust even under serum-free conditions, indicating that TRPC6 activation can promote collagenolytic enzyme release in multiple extracellular contexts.

Collectively, these data demonstrate that TRPC6 activation elicits a selective and temporally structured matrix remodeling response in IVD cells. This response is characterized by sustained, high-magnitude induction of MMP-1 and a delayed but pronounced upregulation of MMP-3, consistent with the establishment of a collagenolytic and proteolytic microenvironment. In contrast, TRPC6 activation does not induce aggrecan-degrading enzymes but instead produces time- and serum-dependent modulation of aggrecan-associated transcription, including a subtle, delayed reduction in ACAN expression and sustained repression of ADAMTS family members under serum-free conditions. Together, these findings indicate that TRPC6 signaling contributes to context-dependent ECM remodeling processes implicated in intervertebral disc degeneration, rather than a generalized or uniform catabolic program.

### 3.6. Hyp9-mediated TRPC6 Activation Induces Angiogenic Signaling and Selectively Modulates Neurotrophic Mediators in Human IVD Cells

The pathological ingrowth of nociceptive nerve fibers (neo-innervation) and blood vessels (neo-vascularization) into the normally avascular and aneural IVD are central features of discogenic pain and advanced degeneration [13, 14]. To determine if TRPC6 signaling contributes to this structural pathology, we next examined whether TRPC6 activation shapes neuro-angiogenic signaling in IVD cells following Hyp9 treatment (1 µM).

Under serum-free conditions, 18 h Hyp9-mediated TRPC6 activation elicited a distinct and selective neuro-angiogenic profile (Figure 6A). The angiogenic factor VEGF was robustly and significantly upregulated, 4.53 ± 0.52-fold (p < 0.0001), exhibiting the most consistent response across donors. NGF expression varied substantially across donors, with increases observed in a subset of samples but not consistently across the cohort, resulting in a non-significant mean increase (3.18 ± 1.00-fold; p = 0.36). In contrast, BDNF (Brain-Derived Neurotrophic Factor) remained unchanged or was slightly downregulated (0.81 ± 0.11-fold; p = 0.12), indicating that TRPC6 signaling selectively engages specific neurotrophic pathways rather than elicits a generalized neurogenic response.

**Figure 6.**
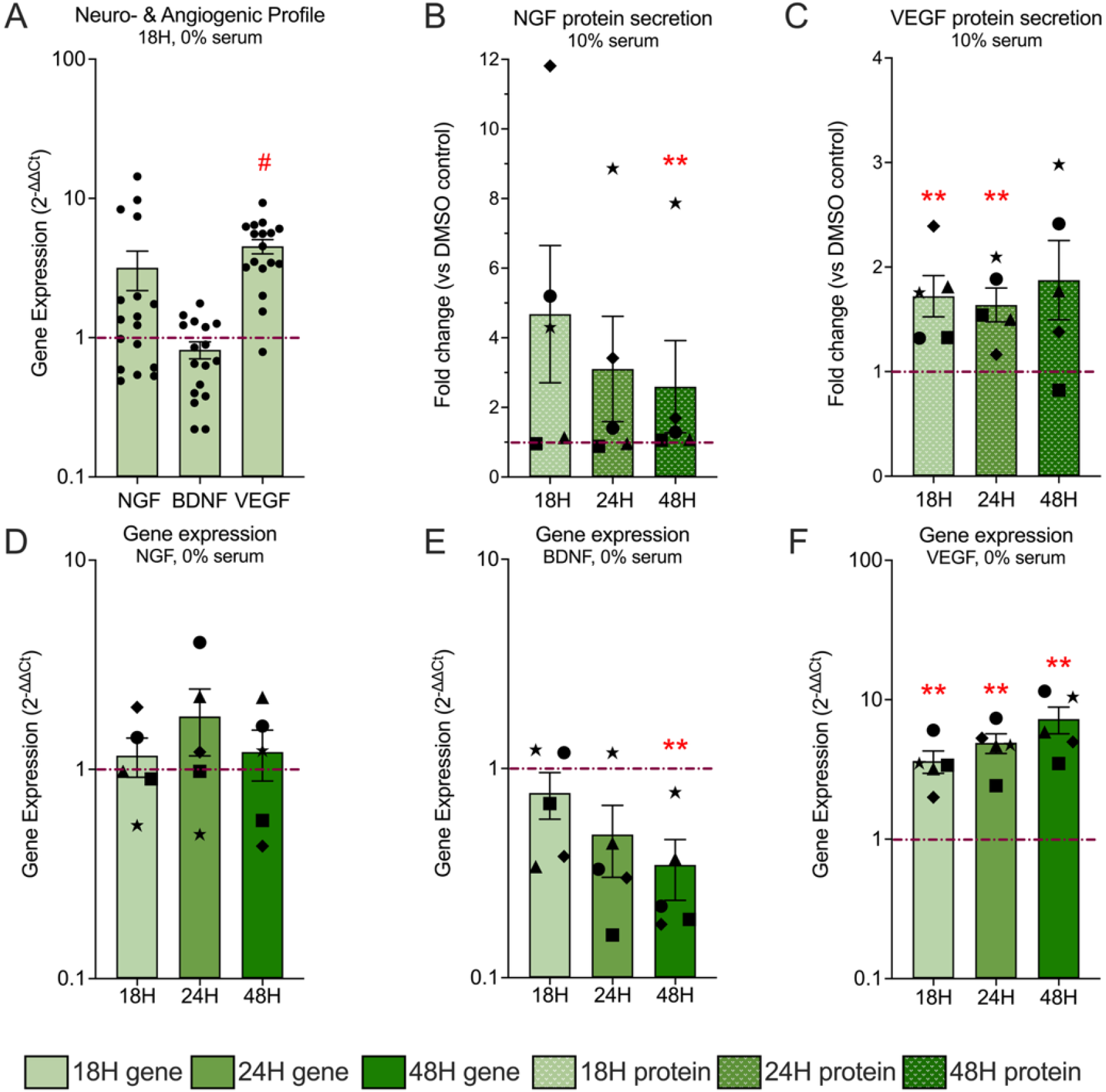
TRPC6 activation modestly modulates neurogenic and angiogenic mediators in human IVD cells. (A) Transcriptional profiling of neuro-angiogenic markers after 18 h of Hyp9 (1 µM) treatment. NGF, BDNF, and VEGF mRNA levels are presented as fold change (2^−ΔΔCt^) relative to vehicle control (dashed line, n= 17). (B-C) Time-dependent secretion of NGF (B) and VEGF (C) protein into the culture media following Hyp9 treatment, quantified by ELISA under serum-replete (10%) conditions and normalized to total DNA content, presented as fold change relative to vehicle control (dashed line, n=5). (D-F) Longitudinal gene expression analysis of NGF (D), BDNF (E), and VEGF (F) over 18, 24, and 48 h, illustrating distinct temporal regulation patterns among neurotrophic and angiogenic mediators (n=5, symbols matched with protein secretion). Data are presented as mean ± SEM with individual donor points; n denotes biological replicates. Statistical significance (**p<0.01, #p<0.0001 vs. vehicle control) was determined by the Mann-Whitney test compared to vehicle control.

Longitudinal analysis (18–48 h) highlighted the distinct kinetics of these factors (Figure 6 D-F). VEGF expression displayed a sustained and progressive increase, rising from 3.63 ± 0.66-fold (p = 0.008) at 18 h to 4.90 ± 0.79-fold (p = 0.008) at 24 h and reaching 7.26 ± 1.57-fold (p = 0.008) at 48 h. This trend was consistent across all donors, suggesting a potent drive for vascularization. NGF expression showed pronounced inter-donor variability over 18–48 h, with a subset of donors exhibiting increased expression at individual time points, whereas others showed little or no change. Consequently, although mean fold changes differed at 18 h (1.16 ± 0.24-fold), 24 h (1.78 ± 0.63-fold), and 48 h (1.21 ± 0.33-fold), none of these changes reached statistical significance (p = 0.68 at all time points). BDNF expression showed progressive reduction over time, a distinct pattern compared with NGF and VEGF. BDNF expression exhibited an early trend toward downregulation that became statistically significant at 48 h time point. BDNF mRNA levels were reduced relative to vehicle at 18 h (0.76 ± 0.19-fold; p = 0.68) and 24 h (0.48 ± 0.18-fold; p = 0.12), reaching a significant decrease at 48 h (0.34 ± 0.11-fold; p = 0.008) (Figure 6B).

To determine whether these transcriptional changes translated to protein secretion, NGF and VEGF release was quantified under 10% serum conditions (Figure 6B–C). VEGF secretion demonstrated a stable and consistently elevated pattern, increasing to 1.72 ± 0.19-fold (p = 0.008) at 18 h and remaining elevated at 24 h (1.63 ± 0.16-fold; p = 0.008) and 48 h (1.87 ± 0.37-fold; p = 0.12). Interestingly, NGF secretion exhibited a heterogeneous temporal pattern following Hyp9 treatment. Although mean protein levels were elevated at earlier time points (18 h: 4.68 ± 1.97-fold; 24 h: 3.10 ± 1.51-fold), these changes were not statistically significant. A modest but significant increase was detected at 48 h (2.59 ± 1.32-fold; p = 0.008). Supplementary confirmation in varying serum conditions (Supplementary Figure S4) corroborated these findings. Notably, ELISA analysis under serum-free conditions confirmed that acute NGF secretion and sustained VEGF release are intrinsic responses to TRPC6 activation, independent of serum factors.

In combination, these data demonstrate that TRPC6 activation drives a specific phenotype characterized by the sustained promotion of angiogenesis (VEGF) alongside variable, context-dependent modulation of NGF, while suppressing BDNF expression. This selective profile aligns with molecular processes associated with nerve ingrowth and vascular infiltration, suggesting that TRPC6 activation may contribute to the development of a microenvironment permissive to neurovascular invasion in degenerating IVDs.

### 3.7. TRPC6-mediated Calcium Influx Recruits MAPK and NF-κB Signaling Pathways in Human IVD Cells

Having established that TRPC6 activation is accompanied by inflammatory, catabolic, and neuro-angiogenic phenotype and given that MAPK and NF-κB pathways are canonical regulators of COX-2, MMP-1, and cytokine expression in the IVD [55], we investigated whether acute TRPC6 activation triggers phosphorylation of these classical intracellular signaling pathway components ERK1/2, p38 and degradation of IκB-α and nuclear translocation of p65.

We first evaluated the phosphorylation status of key MAPKs. Western blot analysis revealed that Hyp9 treatment (1 µM) triggered rapid phosphorylation of ERK1/2 (p44/42) and p38 MAPK within 15 minutes, comparable to the positive control TNF-α (Figure 7A-B). Quantification confirmed an upregulation of kinase activity; phospho-ERK1/2 levels increased to 2.51 ± 1.20-fold (p = 0.31) at 15 minutes and significantly elevated to 2.66 ± 0.66-fold (p = 0.029) at 30 minutes (Figure 7D). The p38 pathway showed a robust and significantly pronounced response, with phospho-p38 levels surging to 4.65 ± 0.48-fold (p = 0.029) at 15 minutes and sustaining a 4.81 ± 1.24-fold (p = 0.029) increase at 30 minutes (Figure 7E). These findings indicate that TRPC6-mediated calcium influx rapidly engages stress-activated MAPK signaling, with particularly strong activation of the p38 pathway.

**Figure 7.**
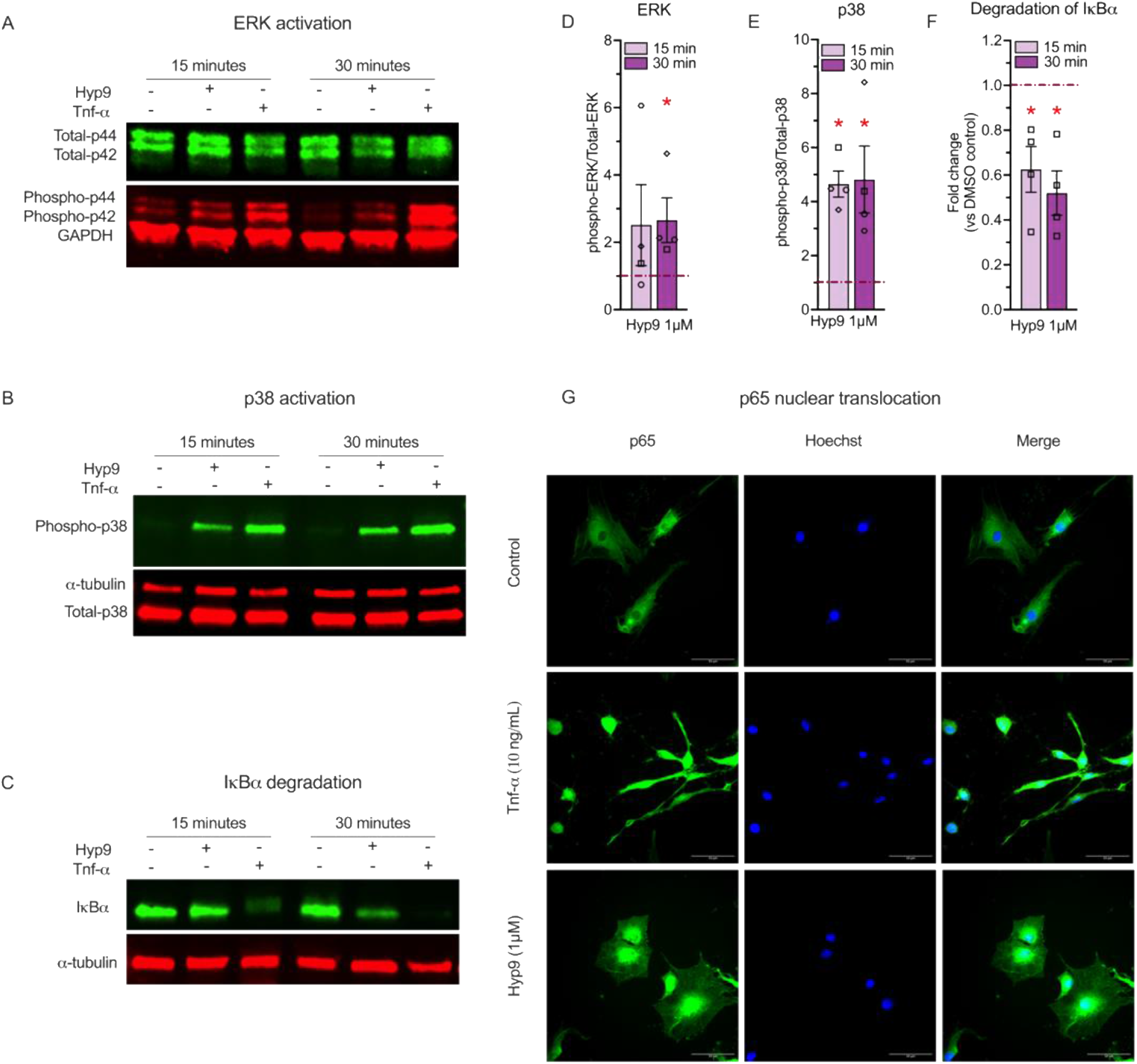
TRPC6 activation recruits MAPK and NF-κB signaling pathways in human IVD cells. Human IVD cells were serum-starved and treated with Hyp9 (1 µM) or the positive control TNF-α (10 ng/mL) for 15 or 30 minutes. (A, D) Representative Western blot and densitometric quantification of ERK1/2 activation. (B, E) Representative Western blot and quantification of p38 MAPK activation, showing a robust increase in phosphorylation (p-p38/Total-p38). (C, F) Representative Western blot images and densitometric analysis of the NF-κB inhibitor IκB-α, showing rapid degradation following stimulation. (G) Representative immunofluorescence images showing the nuclear translocation of the NF-κB subunit p65 (green) following stimulation (Scale bar = 50 µm). Data are presented as fold change relative to vehicle control (dashed line); mean ± SEM, n=4; biological replicates. Statistical significance (*p < 0.05 vs. vehicle control) was determined by Mann-Whitney test compared to vehicle control.

To evaluate if TRPC6 also activates the NF-κB signaling axis, the master regulator of inflammation, we monitored the stability of the inhibitory protein IκB-α. Under basal conditions, IκB-α sequesters the NF-κB complex in the cytoplasm; its degradation is a prerequisite for pathway activation. Hyp9 stimulation resulted in a significant reduction in IκB-α levels (Figure 7C). Quantification revealed that IκB-α abundance dropped to 0.62 ± 0.1-fold (∼38% reduction; p = 0.029) at 15 minutes and persisted at 0.52 ± 0.09-fold (∼48% reduction; p = 0.029) at 30 minutes, mirroring the degradation profile observed with TNF-α (Figure 7F). This rapid loss of IκB-α is consistent with NF-κB pathway activation. To confirm that this degradation is consistent with transcriptional activation, we visualized the localization of the NF-κB subunit p65 using immunofluorescence (Figure 7G). In vehicle-treated cells, p65 localization was predominantly cytoplasmic, whereas Hyp9 stimulation promoted clear nuclear translocation of p65. This redistribution confirms that TRPC6 activation is sufficient to trigger NF-κB nuclear signaling competent for transcriptional regulation.

Together, these findings demonstrate that Hyp9-mediated activation of TRPC6 rapidly recruits inflammatory signaling pathways in IVD cells, including robust activation of p38 MAPK, delayed activation of ERK1/2, and canonical NF-κB signaling marked by IκB-α degradation and p65 nuclear translocation. Given the established roles of the MAPK and NF-κB pathways in regulating inflammatory and catabolic gene expression in the intervertebral disc, their engagement downstream of TRPC6 provides a mechanistic context for the transcriptional responses observed in this study; however, direct pathway dependence will require future inhibition-based analyses.

## 4. Discussion

IVD degeneration is a multifactorial process involving inflammatory cytokine production, ECM breakdown, neurovascular ingrowth, and altered mechanotransduction [56, 57]. Although numerous downstream effectors of degeneration have been extensively described, the upstream signaling events that initiate and coordinate these pathological responses remain incompletely defined. This study identifies TRPC6 as an upstream regulator that simultaneously activates inflammatory, catabolic, and neurovascular pathways in human degenerative IVD cells. Using pharmacological activation of TRPC6, we demonstrate that TRPC6 activation elicits rapid calcium influx, recruits MAPK and NF-κB signaling, and induces a broad transcriptional and secretory response that recapitulates key molecular features of disc degeneration and discogenic pain.

Pharmacological activation of TRPC6 in the present study was achieved using Hyp9, a synthetic derivative of hyperforin that has been widely employed across multiple cellular systems to interrogate TRPC6-focused Ca^2^+ signaling. Among available TRPC6 activators, Hyp9 was selected based on its comparatively well-characterized use and reported engagement of TRPC6 over closely related TRPC isoforms, particularly in non-neuronal cell types [58-60].

TRPC6 is a non-selective cation channel with high permeability to Ca^2^+, and Ca^2^+ influx through TRPC6 is a well-established trigger for downstream MAPK, NF-κB, and transcriptional signaling cascades. TRPC6 has increasingly been recognized as a mechano- and chemo-sensitive ion channel that transduces extracellular stimuli via intracellular calcium signaling to cytoskeletal remodeling, and transcriptional regulation [15, 61]. Consistent with this, the present study focused on Ca^2+^-dependent signaling pathways and transcriptional programs engaged following Hyp9 treatment. Although the observed Ca^2+^ responses are consistent with TRPC6-associated Ca^2+^ entry, contributions from intracellular Ca^2+^ stores, including IP_3_ receptor–mediated endoplasmic reticulum release, cannot be excluded and warrant further investigation. In the IVD, our previous work [45] was the first to demonstrate that TRPC6 expression is closely linked to degeneration severity. Using quantitative gene expression analysis across a large cohort of symptomatic surgical specimens, Sadowska et al. showed that TRPC6 mRNA levels increase progressively from healthy to severely degenerated discs. Complementary immunohistochemical staining performed on an independent tissue set (including asymptomatic and symptomatic degenerated discs obtained as pathology specimens) further demonstrated elevated TRPC6 protein expression in painful degenerated discs compared with asymptomatic degenerative cases. These findings established that TRPC6 upregulation is associated not only with structural tissue degeneration but also with clinically relevant symptomatic states. Building on this, Aripaka et al. (2022) [46] reported a positive correlation between TRPC6 expression and patient-reported visual analogue scale (VAS) pain scores in lumbar disc samples. This independent observation reinforces the notion that TRPC6 is linked to discogenic pain pathways and suggests that heightened TRPC6 activity may contribute to nociceptive sensitization within the degenerating disc. Together, these studies position TRPC6 as a potential upstream mediator of inflammatory and pain-related signaling in human disc pathology.

Gene expression analysis confirmed that TRPC6 was expressed in cells from all human donors included in this study. The isolated human IVD cells were derived from degenerated tissue (Pfirrmann Grade III–V), reflecting a pathological environment characterized by chronic matrix disruption and altered mechanical loading [62, 63]. Cells from such advanced degenerative states exhibit dysregulated mechanosensitivity and altered intracellular signaling, which are important contextual factors when interpreting TRPC6-dependent responses.

Prolonged Hyp9 stimulation reduced TRPC6 transcript levels while total TRPC6 protein abundance remained unchanged, consistent with activity-dependent transcriptional feedback and post-transcriptional buffering described for ion channels [41, 64, 65]. Importantly, the maintenance of TRPC6 protein levels indicates that Hyp9 activates TRPC6 signaling without acutely altering channel availability, supporting the interpretation that downstream inflammatory and catabolic responses arise from functional channel activation rather than changes in expression. These findings also highlight that TRPC6 expression and TRPC6 activation are not necessarily coupled. Although TRPC6 expression has been reported to increase with disc degeneration, this regulation is likely context dependent and not solely driven by channel activity. Indeed, inflammatory stimulation has been shown in other TRP channel contexts to downregulate TRPC6 expression, suggesting that factors beyond inflammation or acute activation contribute to the elevated TRPC6 levels observed in degenerative disc tissue [66]. The chronic, heterogeneous mechanical and biochemical environment of the degenerating disc therefore likely engages regulatory mechanisms distinct from sustained, uniform pharmacological activation in vitro.

One of the most striking findings of this study is the robust and coordinated inflammatory response elicited by Hyp9-mediated TRPC6 activation in human degenerative IVD cells. Among the inflammatory mediators examined, COX-2 (PTGS2) exhibited particularly strong and persistent induction, with expression levels exceeding 100-fold in some donors. The magnitude of this response far exceeded that of other inflammatory mediators, suggesting that TRPC6 activation may be a major upstream driver of prostaglandin synthesis in the degenerative disc environment. Importantly, these cells were derived from Pfirrmann grade III–V tissue, a pathological state characterized by chronic inflammation, matrix disruption, and altered mechanosensitivity, which may sensitize TRPC6 signaling pathways and amplify downstream inflammatory and catabolic responses compared to healthy disc cells [51, 67]. Recognizing this disease-specific context is essential for interpreting the strength of the observed effects and for reconciling these findings with studies employing different cell sources or mechanical environments. COX-2-derived prostaglandins, such as PGE_2_, have extensive roles in nociception and inflammation and are widely implicated in low back pain [68, 69]. Accordingly, the pronounced COX-2 response to TRPC6 activation provides a plausible mechanistic link between TRPC6 signaling, inflammatory amplification, and pain sensitization in degenerative disc disease.

In addition to COX-2, TRPC6 activation also induced IL-6 and IL-8, two cytokines with well-established roles in disc degeneration. IL-6 contributes to inflammatory signaling, matrix remodeling, and chemotaxis, while IL-8 recruits and activates neutrophils and is associated with painful disc herniation [70]. Notably, the sustained induction of IL-8 observed in our study aligns with cytokine profiles reported in degenerated human IVDs ex vivo, in which IL-8 is elevated in a temporally dependent manner, whereas IL-6 expression is often more variable across donors [51, 71-73]. These findings support a model in which TRPC6 activation not only triggers inflammation but also contributes to the chronic, self-amplifying cytokine loop characteristic of advanced disc degeneration.

Our findings also highlight the therapeutic potential of targeting TRPC6. Because TRPC6 sits upstream of multiple inflammatory effectors, its inhibition could simultaneously attenuate several key cytokines and mediators, including COX-2, IL-6, and IL-8. This contrasts with current biologic approaches that act on single downstream inflammatory targets. For example, TNF-α inhibitors have shown limited benefit in discogenic pain [74, 75]. These clinical outcomes likely reflect the complex, multifactorial nature of disc inflammation rather than shortcomings of any individual cytokine pathway. In this context, targeting an upstream regulator of calcium-dependent inflammatory signaling, such as TRPC6, may represent a more effective therapeutic strategy than individual cytokine blockade by simultaneously modulating multiple inflammatory, catabolic, and neurovascular pathways. Our findings support this concept by demonstrating that TRPC6 activation engages coordinated downstream signaling networks rather than isolated cytokine responses. This interpretation is consistent with the neuroimmune framework of discogenic pain described by Cheng et al., [76], which emphasizes the integration of inflammatory, angiogenic, and nociceptive signaling rather than reliance on single-mediator pathways.

A critical feature of disc degeneration is progressive ECM degradation, driven largely by MMPs [54, 77]. In the present study, Hyp9-mediated TRPC6 activation elicited a selective and temporally structured collagenolytic response, characterized by robust MMP-1 upregulation and a temporally delayed but pronounced induction of MMP-3. MMP-1 functions as an initiating collagenase, whereas MMP-3 exhibits broader substrate specificity and can amplify matrix remodeling by activating additional MMPs. The temporal hierarchy observed in our study, early MMP-1 induction followed by delayed MMP-3 activation, suggests that TRPC6 signaling may coordinate sequential phases of collagen degradation rather than triggering a global catabolic response. This pattern aligns with MMP profiles reported in degenerative human discs and supports the role of TRPC6 activation in accelerating matrix deterioration [77].

In contrast, MMP-2 expression was consistently suppressed following TRPC6 activation, highlighting pathway specificity within the MMP family. MMP-2 activity in disc tissue is often regulated post-transcriptionally and depends on membrane-type MMPs, particularly MMP-14 (MT1-MMP), for pro-enzyme activation at the cell surface [78, 79]. The transcriptional suppression of MMP-2 observed here suggests that TRPC6-driven remodeling may preferentially engage collagenase-dominated pathways rather than gelatinase-dependent matrix turnover, potentially shifting the balance of ECM degradation mechanisms in degenerated discs [11, 53].

Analysis of proteoglycan-associated genes further revealed non-uniform regulation of aggrecan remodeling pathways. While ACAN expression was largely preserved at early time points and displayed a minor decline only at later stages, both ADAMTS4 and ADAMTS5 were significantly suppressed, with ADAMTS5 showing sustained repression across all time points examined. These findings indicate that TRPC6 activation does not robustly induce transcription of aggrecanase-related genes and instead is associated with a temporally distinct transcriptional profile, characterized by early induction of collagenolytic enzymes and only modest, delayed changes in aggrecan-associated gene expression. The modest yet significant late reduction in ACAN expression may reflect secondary transcriptional adaptations downstream of sustained TRPC6 signaling or altered cellular homeostatic states, rather than direct activation of aggrecan-degrading enzymes. Together, these data suggest that TRPC6 activation preferentially drives collagen-focused matrix remodeling while differentially modulating proteoglycan-associated pathways, reinforcing the concept of selective, context-dependent ECM remodeling in disc degeneration.

ECM degradation can further promote inflammation through the release of matrix-derived damage-associated molecular patterns (DAMPs), which activate Toll-like receptors such as TLR2 and TLR4, which are highly expressed in degenerative discs and converge on NF-κB signaling [80-82]. Although DAMP release was not directly assessed in this study, the induction of MMPs by TRPC6 suggests that TRPC6 activation contributes to a feed-forward loop of matrix breakdown and inflammatory amplification.

A hallmark of disc degeneration that directly contributes to pain is nerve and blood vessel ingrowth into the normally aneural and avascular disc. In this study, TRPC6 activation consistently induced angiogenic signaling, evidenced by robust and sustained VEGF expression and secretion. VEGF is a central mediator of neovascularization and has been closely linked to degeneration severity, local hypoxia, and vascular infiltration of the disc [13]. Our findings indicate that NGF responses to TRPC6 activation are highly donor-dependent, with variable transcriptional regulation across samples but with measurable accumulation of NGF protein over time. Differences between NGF transcript and protein responses across serum conditions (Supplementary Figure S4) likely reflect context-dependent regulation of NGF at multiple levels. While serum deprivation appeared to accentuate transcriptional variability, NGF protein accumulation was more consistently detected under serum-free conditions, suggesting complex regulation of NGF translation, secretion, and/or extracellular stability downstream of TRPC6 activation. NGF is a potent driver of nociceptor sensitization and is sufficient to induce pain behaviors in animal models; anti-NGF therapies reduce pain in multiple models of musculoskeletal injury, including disc injury [14]. Elevated VEGF and modest NGF in response to TRPC6 activation suggest that TRPC6 may contribute not only to inflammation and matrix breakdown but also to pathological innervation and vascularization, thereby linking TRPC6 activation more directly to discogenic pain.

In contrast to NGF and VEGF, BDNF did not show transcriptional induction following TRPC6 activation and instead exhibited a delayed downregulation. Given that NGF and VEGF are commonly regulated by MAPK-dependent pathways [52, 55, 67, 83], whereas BDNF is more closely associated with CREB-dependent neuronal activity pathways [84, 85], these findings suggest that TRPC6 activation selectively engages inflammatory and angiogenic signaling modules rather than broadly activating neurotrophic programs. Consistent with this interpretation, VEGF induction in multiple tissues has been shown to occur through TRPC6-dependent activation of hypoxia-inducible factor-1 alpha (HIF-1α), nuclear factor of activated T cells (NFAT), and mitogen-activated protein kinase kinase (MEK) pathways that we also observed here, with robust MAPK phosphorylation [86, 87]. Together, the induction of VEGF, selective modulation of NGF, and suppression of BDNF position TRPC6 as an upstream regulator of neurovascular remodeling in the degenerative disc, capable of shaping a microenvironment permissive for pain signaling and vascular infiltration.

As mentioned above, TRPC6 activation led to rapid phosphorylation of p38 and ERK1/2, two MAPKs central to inflammatory signaling, matrix regulation, and nociception. p38 MAPK activation is strongly associated with IL-6 and COX-2 induction in disc cells, while ERK1/2 contributes to MMP production and NGF expression [51, 52, 67, 88]. The activation profiles we observed align with the temporal changes in cytokine and MMP expression, supporting the interpretation that TRPC6-mediated MAPK activation contributes to the inflammatory and catabolic gene programs that follow. In future work, we plan to investigate the involvement of these downstream pathways more directly, including through targeted loss-of-function approaches. In parallel, TRPC6 activation triggered robust NF-κB pathway activation, evidenced by IκB-α degradation and p65 nuclear translocation. NF-κB is a central transcriptional regulator of inflammatory gene expression in the disc, orchestrating cytokine production, adhesion molecule expression, and anti-apoptotic signaling. The rapid activation of NF-κB upon TRPC6 stimulation suggests that TRPC6 lies upstream of this major transcriptional hub, reinforcing its role as a central initiator of inflammatory signaling in degenerative human IVD cells. Taken together, the simultaneous activation of MAPK and NF-κB signaling by TRPC6 provides a mechanistic framework for understanding how TRPC6 activation leads to the coordinated induction of inflammatory, catabolic, and neurovascular mediators observed in this study.

At first glance, our findings appear to differ from those of Shao et al. [89], who reported that TRPC6 activation in human NP cells attenuated stiffness-induced glycolytic and inflammatory signaling. Variations in experimental context can largely reconcile these differences. Shao et al. examined NP-enriched cells derived from moderately degenerated discs, whereas our study used mixed NP/AF populations exclusively from advanced Pfirrmann grade III–V surgical specimens. Cells from advanced degeneration exhibit higher basal inflammation and elevated TRPC6 expression, potentially predisposing them to stronger inflammatory responses upon TRPC6 activation. Moreover, the initiating stimuli differed between studies: TRPC6 activation in Shao et al. occurred in the context of mechanical stress, whereas in our study TRPC6 activation itself served as the primary perturbation. Together, these findings support a context-dependent model in which TRPC6 signaling exerts distinct effects depending on degeneration state, cellular phenotype, and the dominant environmental cues.

Several limitations should be acknowledged. First, our experiments were conducted in a 2D monolayer culture, which does not reproduce the osmotic, mechanical, or three-dimensional architecture of the native disc. Although 2D culture remains a widely used model for signal transduction studies, it may influence the cell phenotype and expression profile. Future studies using 3D culture systems (potentially with mechanical loading bioreactors) will be necessary to evaluate TRPC6 signaling under more physiologically relevant conditions. Second, cell populations derived from degenerative discs often exhibit partial mixing of NP and AF phenotypes due to the loss of tissue boundaries during degeneration. While this mixed-cell environment reflects the in vivo condition of advanced degeneration, it limits the ability to assign TRPC6 responses to specific disc cell types. Third, as with all pharmacological tools, Hyp9-based activation cannot fully exclude contributions from other TRP channels expressed in human IVD cells. Future studies incorporating TRPC6 knockdown or loss-of-function approaches will be important to further refine mechanistic specificity. Lastly, although we identified strong transcriptional and secretory responses to TRPC6 activation, we did not directly measure functional outcomes ex vivo or in vivo. However, this is planned for future studies.

Despite these limitations, our findings have important translational implications. The ability of TRPC6 activation to simultaneously induce inflammatory cytokines, prostaglandin-synthesizing enzymes, matrix-degrading enzymes, and neurovascular mediators suggests that TRPC6 lies upstream of many pathogenic processes in disc degeneration. Because this upstream positioning suggests a therapeutic advantage over approaches targeting individual cytokines or enzymes, we are currently conducting in vitro studies using selective TRPC6 inhibitors to determine whether TRPC6 blockade can reverse or prevent the inflammatory and catabolic programs initiated by Hyp9. In parallel, we are planning in vivo studies that incorporate behavioral pain assessments to evaluate the functional consequences of TRPC6 modulation. Together, these approaches will help determine whether TRPC6 represents a viable therapeutic target for preventing or mitigating discogenic pain and disc degeneration.

## 5. Conclusions

In summary, this study demonstrates that TRPC6 activation is sufficient to induce a comprehensive degenerative phenotype in human IVD cells, engaging inflammatory, catabolic, and neurovascular pathways and it recruits MAPK and NF-κB signaling. Our findings, integrated with existing literature, support a context-dependent role for TRPC6 in disc biology and highlight TRPC6 as a promising upstream therapeutic target. Future in vivo studies will further elucidate the role of TRPC6 in disc health and disease and inform the development of TRPC6-targeted interventions for discogenic pain.

## Supplementary Materials

The following supporting information is available online: Figure S1: Basal TRPC6 expression and donor-specific associations in human intervertebral disc cells; Figure S2: Inflammatory gene expression and protein secretion in human intervertebral disc cells following Hyp9-mediated TRPC6 activation; Figure S3: Catabolic gene expression and protein secretion in human intervertebral disc cells following Hyp9-mediated TRPC6 activation; Figure S4: Neuro-angiogenic gene expression and protein secretion in human intervertebral disc cells following Hyp9-mediated TRPC6 activation; Table S1: Human intervertebral disc donor characteristics; Table S2: TaqMan gene expression assays used for qPCR analysis; Table S3: Primary and secondary antibodies used for Western blot pathway analysis; Table S4: Spearman correlations between basal TRPC6 expression and Hyp9-responsive gene expression.

## Author Contributions

J.V.B. designed and performed the experiments, analyzed the data, prepared the figures, and wrote the original draft. A.M. and V.P. provided human intervertebral disc tissue samples and clinical support. J.P.J., K.Y., and M.E. assisted in disc sample collection and shipment. M.S. assisted with the formulation and interpretation of calcium kinetics analysis. L.S.S. contributed to the interpretation of neuro-inflammatory relevance and provided critical feedback on the manuscript. W.H. provided statistical expertise and assisted with data interpretation. K.W.K. conceived and supervised the project, acquired funding, provided resources, and contributed to manuscript writing and critical revision. All authors have read and agreed to the published version of the manuscript.

## Funding

This study was supported by Eurospine, The Spine Society of Europe (Grant No. 2023-027).

Institutional Review Board Statement: Human intervertebral disc tissue specimens were obtained in de-identified form from clinical partner institutions under protocols approved by their respective Institutional Review Boards (URMC = STUDY00005200; Medstar = STUDY00008525). Because all samples were fully de-identified prior to receipt, the study was determined not to constitute human subjects research at the Rochester Institute of Technology.

## Informed Consent Statement

Informed consent was obtained from all donors involved in this study according to IRB-approved guidelines for the use of de-identified surgical waste tissues.

## Data Availability Statement

The data supporting the findings of this study are available within the article and its Supplementary Materials. Additional raw datasets generated during the current study are available via Figshare (DOI: 10.6084/m9.figshare.30899459) and will be made publicly accessible upon publication.

## Acknowledgment

The authors extend their gratitude to Poorya Esmaili for his assistance with imaging and technical support during data acquisition.

## Abbreviations

The following abbreviations are used in this manuscript:

ADAMTS: A Disintegrin And Metalloproteinase with Thrombospondin Motifs
AF: Annulus Fibrosus
AUC: Area Under the Curve
BCA: Bicinchoninic Acid
BSA: Bovine Serum Albumin
Ca^2+^: Calcium ion
COX2: Cyclooxygenase-2
DAG: Diacylglycerol
DCBP: Discogenic Chronic Back Pain
DH: Disc Herniation
DDD: Degenerative Disc Disease
ECM: Extracellular Matrix
ELISA: Enzyme-Linked Immunosorbent Assay
ERK: Extracellular Signal-regulated Kinase
FBS: Fetal Bovine Serum
GPCR: G Protein-Coupled Receptor
HEPES: 4-(2-Hydroxyethyl)-1-piperazineethanesulfonic Acid
IκB-α: Inhibitor of kappa B alpha
IL: Interleukin
IVD: Intervertebral Disc
LDH: Lactate Dehydrogenase
LysoPC: Lysophosphatidylcholine
MAPK: Mitogen-Activated Protein Kinase
MMP: Matrix Metalloproteinase
NF-κB: Nuclear Factor kappa B
NGF: Nerve Growth Factor
NP: Nucleus Pulposus
PBS: Phosphate-Buffered Saline
PLC: Phospholipase C
RIPA: Radioimmunoprecipitation Assay
TNF-α: Tumor Necrosis Factor alpha
TRP Transient: Receptor Potential
TRPC6: Transient Receptor Potential Canonical 6
VEGF: Vascular Endothelial Growth Factor

