## Supplemental information for "TRPC6-Mediated Ca^2+^ Influx Activates MAPK and NFκB Signaling and Elicits Pro-Inflammatory and Catabolic Responses in Human Intervertebral Disc Cells"

Table S1. Human intervertebral disc donor characteristics

| Donor Number | | Level | Pfirrmann Grade | Diagnosis | Sex | BMI | Age | Steroid injection |
|---|---|---|---|---|---|---|---|---|
| 1 | | C4-5 | 4 | Stenosis | Male | 25.1 | 79 | No |
| 2 | | L4-5/ L5-S1 | 4 | DDD | Male | 33.6 | 47 | Yes |
| 3 | | L3-4/ L4-5 | 4 | Stenosis | Female | 30.7 | 75 | Yes |
| 4 | | n.a. | n.a. | n.a. | n.a. | n.a. | n.a. | n.a. |
| 5 | | n.a. | n.a. | n.a. | n.a. | n.a. | n.a. | n.a. |
| 6 | | L4-5 | 5 | DDD | Female | 30 | 45 | n.a. |
| 7 | | L5-S1 | 3 | DH | Male | 23.8 | 50 | Yes |
| 8 | | L5-S1 | 5 | DH | Female | 38.7 | 33 | Yes |
| 9 | | L4-5/ L5-S1 | 4 | DH | Male | 26 | 42 | Yes |
| 10 | | C6-7 | 3 | DH | Female | 30.8 | 54 | No |
| 11 | | L5-S1 | 4 | DH | Male | 22.3 | 27 | Yes |
| 12 | | L5-S1 | 4 | DH | Female | 32.5 | 29 | Yes |
| 13 | ● | L5-S1 | 5 | DH | Male | n.a. | 36 | n.a. |
| 14 | ■ | C5-6 | 3 | DH | Female | n.a. | 40 | n.a. |
| 15 | ▲ | L5-S1 | 4 | DH | Male | n.a. | 41 | n.a. |
| 16 | ◆ | L5-S1 | 3 | DH | Female | 22.3 | 36 | Yes |
| 17 | ★ | L5-S1 | 3 | DH | Female | 20.7 | 42 | Yes |

For samples 4 and 5, detailed Pfirrmann grading information was not available; however, based on clinical assessment, these samples fall within the Pfirrmann grade III–V range.

Abbreviations: n.a., not available (clinical or demographic information not provided for these donors); DDD, Degenerative disc disease; DH, Disc herniation.

Table S2. TaqMan gene expression assays used for qPCR analysis

| **Nr.** | **Target Gene** | **Gene Name** | **Assay ID** |
|---|---|---|---|
| **1** | *TRPC6* | *Transient receptor potential cation channel subfamily C member 6* | Hs00988479_m1 |
| **2** | *IL6* | *Interleukin 6* | Hs00174131_m1 |
| **3** | *CXCL8 (IL8)* | *C-X-C motif chemokine ligand 8 (interleukin 8)* | Hs00174103_m1 |
| **4** | *PTGS2 (COX2)* | *Prostaglandin-endoperoxide synthase 2* | Hs00153133_m1 |
| **5** | *MMP1* | *Matrix metallopeptidase 1* | Hs00899658_m1 |
| **6** | *MMP2* | *Matrix metallopeptidase 2* | Hs01548727_m1 |
| **7** | *MMP3* | *Matrix metallopeptidase 3* | Hs00968305_m1 |
| **8** | *MMP13* | *Matrix metallopeptidase 13* | Hs00942584_m1 |
| **9** | *ACAN* | *Aggrecan* | Hs00153936_m1 |
| **10** | *ADAMTS4* | *ADAM metallopeptidase with thrombospondin type 1 motif 4* | Hs00192708_m1 |
| **11** | *ADAMTS5* | *ADAM metallopeptidase with thrombospondin type 1 motif 5* | Hs01095518_m1 |
| **12** | *BDNF* | *Brain derived neurotrophic factor* | Hs02718934_s1 |
| **13** | *NGF* | *Nerve growth factor* | Hs00171458_m1 |
| **14** | *VEGFA* | *Vascular endothelial growth factor A* | Hs00900055_m1 |
| **15** | *YWHAZ* | *Tyrosine 3-monooxygenase/tryptophan 5-monooxygenase activation protein zeta* | Hs01122445_g1 |

All assays were pre-designed TaqMan Gene Expression Assays (FAM-MGB) for Homo sapiens (Thermo Fisher Scientific). Gene expression was normalized to *YWHAZ* and analyzed using the ΔΔCt method.

Table S3. Primary and secondary antibodies used for Western blot pathway analysis

| **Nr** | **Antibody** | **Dilution** | **Catalog Nr.** | **Company** |
|---|---|---|---|---|
| 1 | $\alpha$-Tubulin (Rabbit) | 1:2000 | 2144 | Cell signaling |
| 2 | GAPDH (Rabbit) | 1:2000 | 2118 | Cell signaling |
| 3 | p44/42 ERK (Mouse) | 1:2000 | 4696 | Cell signaling |
| 4 | Phospho-p44/42 ERK (rabbit) | 1:2000 | 4370 | Cell signaling |
| 5 | p38 MAPK (Rabbit) | 1:2000 | 9212 | Cell signaling |
| 6 | Phospho-p38 MAPK (Mouse) | 1:2000 | 9216 | Cell signaling |
| 7 | Inhibitor of NF-κB alpha (IKB-$\alpha$) (Mouse) | 1:2000 | 4814 | Cell signaling |
| 8 | IRDye® 800CW Goat Anti-Mouse IgG1 | 1:20,000 | 926-32350 | LICORbio |
| 9 | IRDye® 680RD Goat Anti-Rabbit IgG1 | 1:20,000 | 926-68071 | LICORbio |

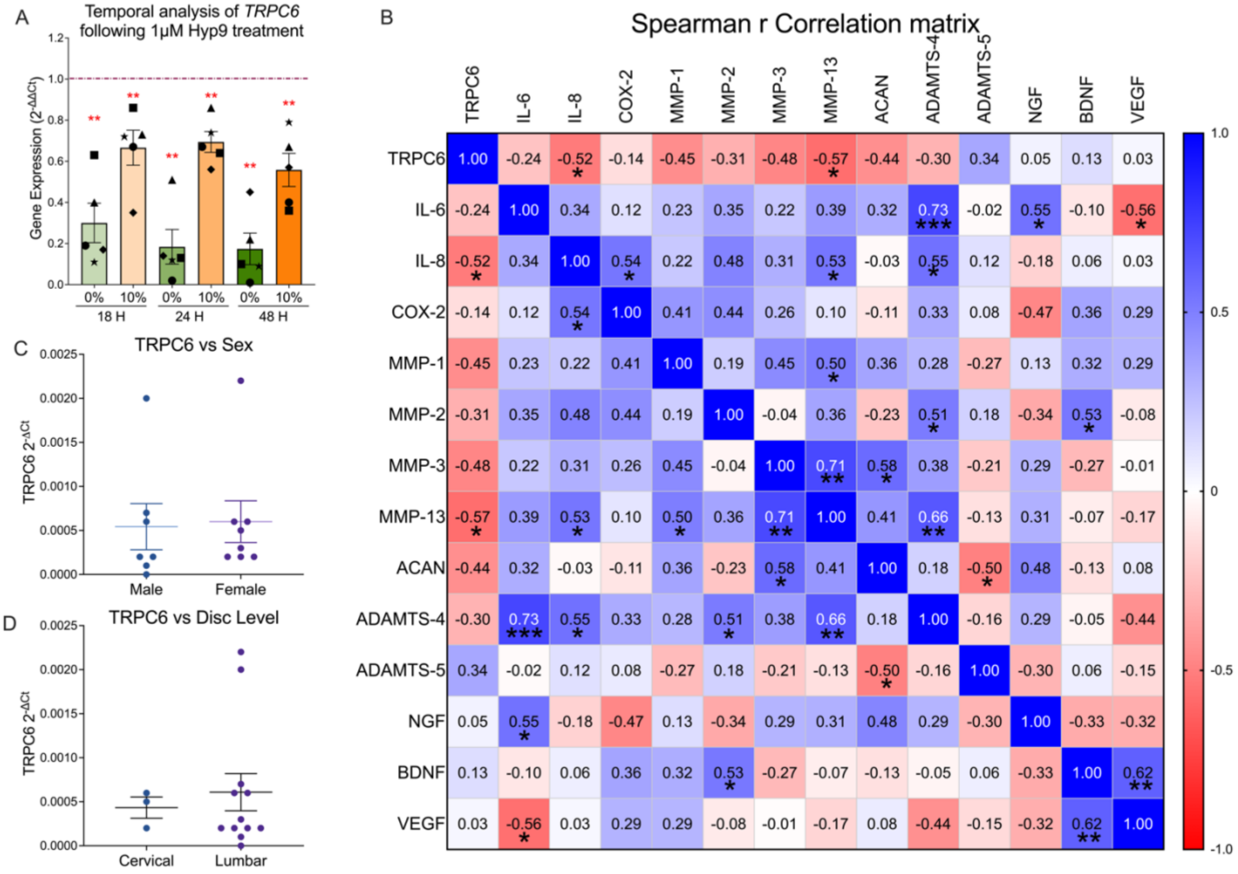


Figure S1. Basal TRPC6 expression and donor-specific associations in human intervertebral disc cells. (A) TRPC6 gene expression under serum-free (0%) and serum-containing (10%) conditions assessed at 18, 24, and 48 h by quantitative PCR following Hyp9 treatment. Data are normalized to YWHAZ and presented as fold change (2^−ΔΔCt) relative to vehicle control (dashed line; n = 5 biological replicates). Statistical significance (**$p < 0.01$ vs. vehicle control) was assessed using the Mann–Whitney test. (B) Spearman correlation matrix showing associations between basal TRPC6 expression and selected inflammatory, catabolic, and neuroangiogenic gene transcripts. Correlation coefficients (r) were calculated using TRPC6 2^−ΔCt values from vehicle control samples (n = 17 donors) and gene expression values from Hyp9-treated samples (1 μM, 18 h, serum-free; n = 17 donors). Correlation strength is represented by the color scale (−1 to +1), with r values displayed within each cell. Statistical significance is indicated as *$p \le 0.05$, **$p \le 0.01$, and ***$p \le 0.001$ (C) TRPC6 expression stratified by sex. Basal TRPC6 $2^{-\Delta Ct}$ values from vehicle control samples in male and female donors are shown as individual data points with mean ± SEM overlaid. Statistical significance was assessed using the Mann–Whitney test. (D) TRPC6 expression across disc regions. Comparison of basal TRPC6 $2^{-\Delta Ct}$ values from vehicle control samples between cervical and lumbar disc regions. Individual donor values are shown with mean ± SEM overlaid. Statistical significance was assessed using the Mann–Whitney test.

Table S4. Spearman correlations between basal TRPC6 expression and Hyp9-responsive gene expression.

| Nr. | TRPC6 Vs Genes (2^- ΔCt) | Spearman r value | p Value |
|---|---|---|---|
| 1 | IL-6 | -0.29 | 0.25 |
| 2 | IL-8 | 0.08 | 0.76 |
| 3 | COX-2 | 0.10 | 0.68 |
| 4 | MMP-1 | -0.09 | 0.73 |
| 5 | MMP-2 | -0.71 | 0.002 |
| 6 | MMP-3 | 0.11 | 0.66 |
| 7 | MMP-13 | -0.08 | 0.76 |
| 8 | ACAN | -0.10 | 0.72 |
| 9 | ADAMTS-4 | -0.68 | 0.003 |
| 10 | ADAMTS-5 | -0.15 | 0.55 |
| 11 | NGF | -0.19 | 0.45 |
| 12 | BDNF | -0.17 | 0.51 |
| 13 | VEGF | -0.30 | 0.22 |

Spearman correlation coefficients (r) were calculated using TRPC6 expression values expressed as 2^−ΔCt (vehicle control) and gene expression values expressed as 2^−ΔΔCt following Hyp9 treatment.

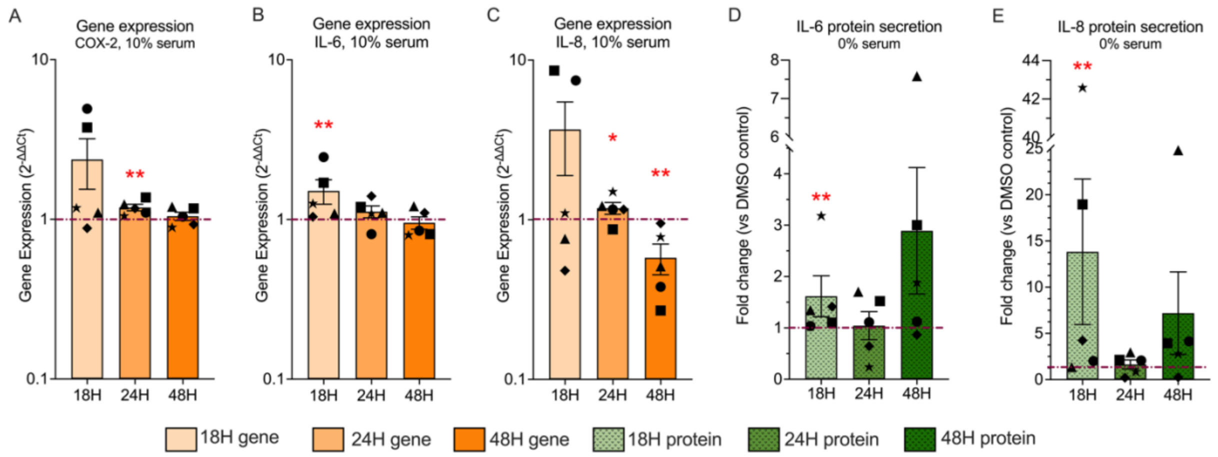


Figure S2. Inflammatory gene expression and protein secretion in human intervertebral disc cells following Hyp9-mediated TRPC6 activation. (A–C) Gene expression of COX-2 (PTGS2) (A), IL-6 (B), and IL-8 (CXCL8) (C) measured at 18, 24, and 48 h under 10% serum conditions by quantitative PCR. mRNA levels are normalized to YWHAZ and presented as fold change (2^−ΔΔCt) relative to vehicle control (dashed line; n = 5). (D–E) Protein secretion of IL-6 (D) and IL-8 (E) measured in conditioned media collected at 18, 24, and 48 h under serum-free (0%) conditions. Protein levels were normalized to total DNA content and are presented as fold change relative to vehicle control (dashed line; n = 5). Data are presented as mean ± SEM (biological replicates). Statistical significance (*p ≤ 0.05, **p ≤ 0.01, #p ≤ 0.0001 vs. vehicle control) was assessed using the Mann–Whitney test.

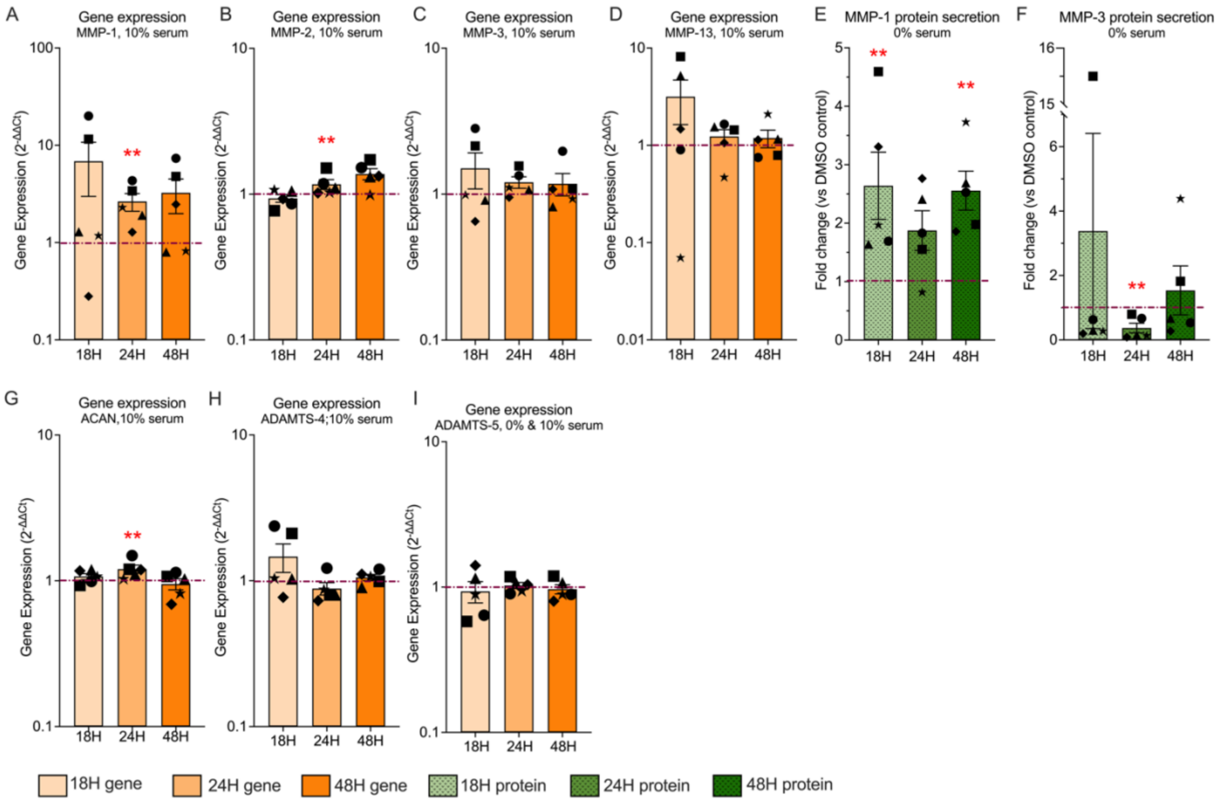


Figure S3. Catabolic gene expression and protein secretion in human intervertebral disc cells following Hyp9-mediated TRPC6 activation. (A–D) Gene expression of MMP-1 (A), MMP-2 (B), MMP-3 (C), and MMP-13 (D) measured at 18, 24, and 48 h under 10% serum conditions by quantitative PCR. mRNA levels are normalized to YWHAZ and presented as fold change (2^−ΔΔCt) relative to vehicle control (dashed line; n = 5). (E–F) Protein secretion of MMP-1 (E) and MMP-3 (F) measured in conditioned media collected at 18, 24, and 48 h under serum-free (0%) conditions. Protein levels were normalized to total DNA content and are presented as fold change relative to vehicle control (dashed line; n = 5). (G–J) Gene expression of ACAN (G) ADAMTS-4 (H), and ADAMTS-5 (I) measured at 18, 24, and 48 h under serum-free (0%) and serum-containing (10%) conditions by quantitative PCR. Data are presented as fold change (2^−ΔΔCt) relative to vehicle control (dashed line; $n = 5$). Data are presented as mean ± SEM (biological replicates). Statistical significance (**$p \leq 0.01$, #$p \leq 0.0001$ vs. vehicle control) was assessed using the Mann–Whitney test.

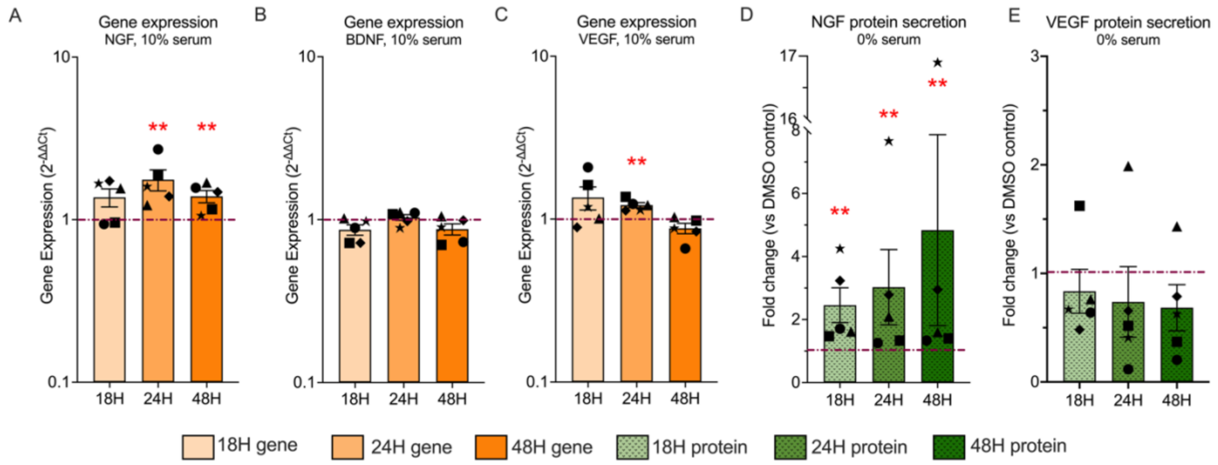


Figure S4. Neuro-angiogenic gene expression and protein secretion in human intervertebral disc cells following Hyp9-mediated TRPC6 activation. (A–C) Gene expression of NGF (A), BDNF (B), and VEGF (C) measured at 18, 24, and 48 h under 10% serum conditions by quantitative PCR. mRNA levels are normalized to YWHAZ and presented as fold change ($2^{-\Delta\Delta Ct}$) relative to vehicle control (dashed line; n = 5). (D–E) Protein secretion of NGF (D) and VEGF (E) measured in conditioned media collected at 18, 24, and 48 h under serum-free (0%) conditions. Protein levels were normalized to total DNA content and are presented as fold change relative to vehicle control (dashed line; n = 5). Data are presented as mean ± SEM (biological replicates). Statistical significance (**$p \leq 0.01$ vs. vehicle control) was assessed using the Mann–Whitney test.
